# A cell-autonomous mechanism regulates BCAA catabolism in white adipocytes and systemic metabolic balance

**DOI:** 10.1101/2023.07.31.551146

**Authors:** Ashley M. Aguillard, Joyce Tzeng, Ismael Ferrer, Bjorn T. Tam, Damaris N. Lorenzo

## Abstract

Elevated plasma branched-chain amino acids (BCAAs) are strongly associated with obesity, insulin resistance (IR), and diabetes in humans and rodent models. However, the mechanisms of BCAA dysregulation and its systemic, organ, and cell-specific implications in the development of obesity and IR are not well understood. To gain mechanistic insight into the causes and effects of plasma BCAA elevations, we leveraged mouse models with high circulating BCAA levels prior to the onset of obesity and IR. Young mice lacking ankyrin-B in white adipose tissue (WAT) or bearing an ankyrin-B variant that causes age-driven metabolic syndrome exhibit downregulation of BCAA catabolism selectively in WAT and excess plasma BCAAs. Using cellular assays, we demonstrated that ankyrin-B promotes the surface localization of the amino acid transporter Asct2 in white adipocytes, and its deficit impairs BCAA uptake. Excess BCAA supplementation worsened glucose tolerance and insulin sensitivity across genotypes. In contrast, BCAA overconsumption only increased adiposity in control mice, implicating WAT utilization of BCAAs in their obesogenic effects. These results shed light into the mechanistic underpinnings of metabolic syndrome caused by ankyrin-B deficits and provide new evidence of the relevance of WAT in the regulation of systemic BCAA levels, adiposity, and glucose homeostasis.

**ARTICLE HIGHLIGHTS:**

- Ankyrin-B deficits in adipose tissue result in elevated circulating BCAAs before the onset of obesity and insulin resistance.
- Ankyrin-B promotes the surface localization of the amino acid transporter Asct2 in white adipocytes and BCAA uptake.
- Excess BCAA supplementation worsens glucose tolerance and insulin sensitivity in ankyrin-B deficient mice.
- BCAA utilization by white adipose tissue is required for the obesogenic effects of BCAA overconsumption.

## Introduction

Branched-chain amino acids (BCAA) (leucine, valine, and isoleucine) are essential amino acids, and as such must be obtained through dietary protein consumption. Thus, circulating levels of BCAA are determined by the combination of their ingested amounts and the rate of their utilization by multiple tissues and organs. Several studies in human cohorts have established the elevated concentration of circulating BCAA and related metabolites as robust biomarkers of cardiometabolic dysregulation, including obesity, insulin resistance (IR), and type II diabetes (T2D). First observed a few decades ago (1, 2), this correlation has been significantly expanded more recently through multiple studies in humans (3-8) and rodents (9-14). Unbiased metabolomics profiling in humans have identified high elevated BCAA as the clearest predictor for developing diabetes in individuals with normal insulin sensitivity (3, 15-17). Correspondingly, BCAA ingestion in healthy humans can impair glucose disposal (18) and dietary BCAA restriction in patients with T2D resulted in improved oral glucose sensitivity (19). Moreover, supplementation of BCAA to a high fat diet (HFD) leads to glucose intolerance in rats (3), while dietary BCAA restriction improved insulin sensitivity in rats and mice (11, 12, 20). Despite these supporting observations, a causal role for BCAAs in IR and the pathogenesis of metabolic syndrome is not clear (21, 22).

The mechanisms of BCAA dysregulation, including its systemic, organ and cell-specific implication in metabolic syndrome, are not well understood. Current models propose a feedforward mechanism where IR can cause BCAA elevation and, conversely, high BCAA levels contribute to IR, particularly in the context of excess lipid availability (21, 22). Tracing studies in mice using stable isotope-labeled BCAAs suggest skeletal muscle as the primary organ of BCAA oxidation (23). However, increased oxidation of BCAA by muscle (or liver, or both) as the result of the muscle-, liver-specific, or combined tissue deletion of the branched-chain ketoacid dehydrogenase kinase *Bckdk*, which inhibits BCAA catabolism by phosphorylating the rate-limiting branched-chain α-ketoacid dehydrogenase *Bckdh*, had no appreciable effect on insulin sensitivity in mice (24). On the other hand, genetic rodent models (9, 10, 25) and human studies (26, 27) of obesity have shown suppression of BCAA catabolic genes in white adipose tissue. Adipose-specific overexpression of the glucose transporter 4 (Glut4) in mice led to down-regulation of BCAA metabolizing enzymes selectively in adipose tissue, elevated circulating BCAA, and systemic IR (9). Transplantation of normal adipose tissue into these mice reduced the plasma BCAA levels (9). Moreover, bariatric surgery-induced weight loss correlates with reduced plasma BCAA and elevation of BCAA catabolic enzymes in adipose tissue (28, 29). These studies underscore the importance of adipose tissue as a major regulator of systemic BCAA levels.

We recently reported that knock-in mice expressing the T2D-associated human ankyrin-B (AnkB) rare variant R1788W, (*Ank2^R1788W/R1788W^;* heretofore AnkB-RW), encoded by *Ank2*, developed metabolic syndrome characterized by primary pancreatic β-cell insufficiency, diet- or age-dependent adiposity, and IR without major changes in appetite or activity (30). In contrast, young (3-month-old) AnkB-RW mice have no significant differences in body weight or composition relative to littermate controls but exhibit increased insulin sensitivity (30). At this age, AnkB-RW mice already display increases in white adipocyte size (30) through an adipose tissue-autonomous mechanism (31) involving increased glucose uptake and de novo lipogenesis due to elevated cell surface localization of Glut4 as the result of impaired Glut4 trafficking. Young AnkB-RW are initially protected against hyperglycemia through the higher glucose uptake by both white adipose tissue (WAT) and skeletal muscle, which compensates for their impaired insulin secretion, but this effect is eventually overridden by the lipotoxic effects of the developed adiposity (30).

In this study, we now report that non-obese young AnkB-RW mice exhibit significant downregulation of BCAA catabolic enzymes selectively in WAT together with excess plasma BCAA, which makes them an attractive model to study effects of BCAA elevation prior to the onset of obesity and diabetes. We show that the elevated circulating BCAA in AnkB-deficient mice is likely driven by impaired BCAA utilization in WAT. Moreover, excess BCAA through diet supplementation failed to increase adiposity in these mice within the context of both a normal and a high fat diet (HFD), but worsened glucose tolerance and insulin sensitivity. Finally, we show that AnkB is required for the proper localization of the amino acid transporter Asct2 at the cell surface of white adipocytes, and its deficit leads to impaired BCAA uptake.

## RESEARCH DESIGN AND METHODS

### Mouse lines and animal care

Experiments were performed in accordance with the guidelines for animal care of the Institutional Animal Care and Use Committee of the University of North Carolina at Chapel Hill and the University of Pennsylvania. Studies were conducted in four-month-old congenic male mice. All mice were housed at 22°C ± 2°C on a 12-hour-light/12-hour-dark cycle and fed *ad libitum* regular chow and water. For diet studies, 5-week-old mice were divided into four groups and fed *ad libitum* either control diet (CD), high-fat diet (HFD) or BCAA-enriched diets (CD-BCAA, HFD-BCAA) for 12 weeks. On a caloric basis, the CD consisted of 10% fat, 20% protein, and 70% carbohydrate (Research Diet D12450K). For the HFD, 45% of the calories were in the form of fat, 20% of the calories were from protein, and 35% of the calories were from carbohydrates (Research Diet D12492). The BCAA enriched diets (CD-BCAA and HFD-BCAA) have the same macronutrient composition as the CD and HFD, respectively. However, half of casein, the main protein source in the CD and HFD formulations, was replaced with individual amino acids, including 150% more BCAA (Research Diet D17011602 and D17050303). See diet composition in Supplemental Material Table 1. Homozygous knock-in mice carrying the p.R1788W variant in the mouse *Ank2* gene (AnkB-RW) and mice with conditional ankyrin-B knockout in adipose tissue (AT-AnkB KO) have been previously reported (30, 31).

### Plasmids

Full-length (FL) 220-kDa AnkB-GFP, MBD AnkB-GFP, and ZU5-Ct AnkB-GFP were previously described (31). To generate the FL mCherry-Asct2 plasmid, we linearized the pmCherry-N1 vector (Takara) by restriction enzyme digestion with FastDigest EcoRI (Thermo Fisher). The coding sequence of human ASCT2 was amplified with homology arms from pDONR221_SLC1A5, a gift from RESOLUTE Consortium & Giulio Superti-Furga (Addgene plasmid #131974; http://n2t.net/addgene:131974; RRID: Addgene_131974) using high fidelity Phusion polymerase (Takara) and primers hSlc1a5_FOR: 5’-TCAAGCTTCGAATTCATGGTGGCCGACCCC-3’ and hSlc1a5_REV: 5’-GGCGACCGGTGGATCCATCACGCTCTCCTTCTCAG-3’. Linearized fragments were assembled using In-Fusion Cloning reagents (Takara). All plasmids were verified by full-length sequencing prior to transfection.

### Antibodies and fluorescent dyes

Affinity-purified rabbit antibodies against AnkB and GFP were generated in our laboratory and have been previously described (30, 31). Other antibodies included mouse anti-alpha 1 sodium potassium ATPase (H-3, sc-48345), mouse anti-GFP (B-2, sc-9996), mouse anti-BCKDE1A (H-5, sc-271538), mouse anti-CD98 (E-5, sc-376815), mouse anti-FASN (G-11, sc-48357), and mouse anti-LAT1 (D-10, sc-374232) from Santa Cruz Biotechnology. Rabbit antibodies against mCherry (ab167453) and BCKDHA (phospho S293, ab200577) were purchased from Abcam. We also used rabbit anti-α-tubulin (11224-1-AP), rabbit anti Fabp4 (12802-1-AP), and mouse anti-GAPDH (2D4, 60004-I-AP) from Proteintech; rabbit anti-ASCT2 (ANT-082) from Alomone; and rabbit antibodies against GAPDH (D16H11 XP) and ACL (4332) from Cell Signaling Technologies. Secondary antibodies used for fluorescence imaging were purchased from Life Technologies and included donkey anti-rabbit IgG conjugated to Alexa Fluor 568 (#A10042) and donkey anti-mouse IgG conjugated to Alexa Fluor 568 (#A10037). Fluorescent signals in western blot analysis were detected using goat anti-rabbit 800CW (926-32211) and goat anti-mouse 680RD (926-68070) from LiCOR. Lipid droplets and nuclei were visualized by BODIPY™ 493/503 (D3922) and DAPI (D1306) staining, respectively, purchased from Life Technologies.

### Body composition and calorimetry analysis

Body composition of 5-month-old control and AnkB-RW male mice was measured by magnetic resonance imaging (MRI). Mice were housed in individual indirect calorimetry cages (Phenomaster/Labmaster, TSE SYSTEMS, Chesterfield, MO) for continuous measurements of O_2_ consumption (VO_2_), CO_2_ production (VCO_2_), activity (total sum of X, Y and Z beam break counts), and food and water consumption for 48 hours. Energy expenditure and respiratory exchange rate [RER (VCO_2_/VO_2_)] were also calculated.

### Tolerance tests

Glucose (OGTT) and insulin (ITT) tolerance tests were performed after 12 weeks of the experimental diet intervention. Mice were fasted for eight hours before the GTT and ITT assays. For GTT evaluation, 2 g/kg body weight of glucose (Sigma) was administered by oral gavage. For ITT, mice were administered 0.75 U/kg body weight of recombinant human insulin (Humulin R; Eli Lilly) by intraperitoneal injection. Blood samples were measured by tail bleeding at 0, 30, 60, 90, and 120 min after glucose or insulin administration and analyzed with an AccuCheck glucometer.

### Amino acid profiling

Amino acid profiles were derived from serum following sample handling and extraction previously described (11). Amino acid profiling was performed by tandem mass spectrometry (MS/MS) using stable-isotope-dilution with internal standards from Isotec, Cambridge Isotopes Laboratories, and CDN Isotopes (11).

### RNA isolation

Total RNA from WAT, BAT, skeletal muscle, and liver was purified using the RNeasy Lipid Tissue Kit (Qiagen) and subjected to on-column DNase treatment (RNase-Free DNase Set, Qiagen) following the manufacturer’s instructions. RNA purity and integrity was assessed using a BioAnalyzer.

### Gene expression profiling from WAT

Gene expression from mouse WAT of 3-month-old control and AnkB-RW mice (n=3/genotype) was profiled using the GeneChip™ Mouse Genome 430 2.0 Array (Affymetrix) Integrative Cancer Genomics Shared Resource at Duke University. Data was analyzed using the Partek Genomics Suite Analysis Software. Pathway analysis was conducted using the Ingenuity Pathway Analysis (Qiagen) software with a cutoff of p<0.05 and a fold change of 2. Interaction network analysis was conducted in Cytoscape.

### Quantitative PCR analysis

cDNA libraries were prepared using the SuperScript IV cDNA synthesis kit (ThermoFisher Scientific). Gene expression was assessed by quantitative PCR using 100 ng of each cDNA and PowerUp™ SYBR™ Green Master Mix detection reagent (ThermoFisher Scientific) in a QuantStudio™ 7 Flex Real-Time PCR system. Primer sequences (listed below) were designed with Primer Express Software (Applied Biosystems) and primers synthesized by IDT DNA.

BCAA catabolism genes: Bcat2 forward 5′-TTCCCAGAGACTCAAGAGAAGTGC-3′ and reverse 5′-GGAGATAGAGAAGAAGCCAAGGTTC-3′; Bchdha forward 5′-AAGCCTCTTCTCCGATGT-3′ and reverse 5′-GCAAAAGTCACCCTGGAATGC-3′; Bckdhb forward 5′-GTGCCCTGGATAACTCATTAGCC-3′ and reverse 5′-CAAACTGGATTTCCGCAATAG-3′; Ivd forward 5′-CATCAGTGGTGAGTTCATCGGAG-3′ and reverse 5′-CAAATCTGTCTTGGCATACACGAC-3′; Dbt forward 5′-GGAAAAGCAGACAGGAGCCATAC-3′ and reverse 5′-TCTCATCACAATACCCGAAGTGG-3′. Amino Acid transporters: Slc1a5 forward 5′-GGTGGTCTTCGCTATCGTCTTTG-3′ and reverse 5′-GCAGGCAGCACAGAATGTATTTG-3′; Slc3a2 forward 5′-TGAGTCCAGCATCTTTCACATCC-3′ and reverse 5′-AGTTGAGCACCACCAGGTAACG -3′; Slc7a5 forward 5′-CTACTTCTTTGGTGTCTGGTGGAA-3′ and reverse 5′-GAGGTACCACCTGCATCAACTTC-3′. GAPDH (primers forward 5′-AGTGCCAGCCTCGTCCCGTA-3′ and reverse 5′-GCCACTGCAAATGGCAGCCC-3′) was used as internal control for normalization.

### Mouse embryonic fibroblast (MEFs) culture and differentiation into adipocytes

Primary MEFs cultures were established from postnatal day 0 (PND0) control and AnkB-RW mice following a described protocol (30, 31). In brief, after removal of all internal organs, bodies were minced and digested in 1X HBSS (Genesee) containing 0.25% trypsin (Life Technologies) and 0.2 mg/ml DNAse (Sigma) for 30 min at 37°C. Tissue was washed three times with 1X HBSS and dissociated in MEF media (Dulbecco’s modified Eagle’s medium (DMEM, Genesee) supplemented with 10% fetal bovine serum (FBS, Genesee), non-essential amino-acids (MEM, Life Technologies), 2 mM glutamine (Life Technologies), and penicillin/streptomycin (Life Technologies), and vigorously agitated for 30 seconds. Dissociated cells were then passed through a 100 µm disposable cell strainer to remove any residual non-dissociated tissue, pelleted by centrifugation at 1,500 rpm for 5 minutes, plated on a 100-mm tissue culture dishes containing MEF media, and incubated at 37°C and 5% of CO_2_ until the culture reached 100% confluence. 48-hours post-confluence MEFs were induced to undergo adipogenic differentiation by incubation for two days with induction media (MEF media supplemented with 1.8 uM insulin (Humulin R; Eli Lilly), 5 nM dexamethasone (Sigma), 10 mM rosiglitazone (Sigma), 0.5 mM 3-isobutyl-1-methylxanthine (IBMX, Cayman). Cells were then switched to maintenance media (MEF media supplemented with 1.8 nM insulin, 10 mM rosiglitazone) for an additional two days. Differentiated adipocytes were kept in MEF media until use.

### *In vitro* BCAA and/or oleic acid supplementation of adipocytes

Differentiated adipocytes were incubated with either MEF media or BCAA-enriched MEF media supplemented with 1.2 mM of each valine, leucine, and isoleucine (Sigma). This results in a final 2 mM concentration of each BCAA in the media and in a 1.5-fold increase in total BCAA content. To evaluate if supplementation with either BCAA, oleic acid, or both enhances adipogenesis, starting at differentiation day 2 adipocytes were incubated with either MEF media, BCAA-enriched MEF media (1.2 mM of each BCAA), oleic acid MEF media (500 µM), or dual BCAA/oleic acid-enriched MEF media.

### Lysine uptake assay

Differentiated adipocytes cultured in 12-well plates were assessed for their ability to uptake lysine nine days after induction of differentiation. In brief, cells were serum starved for 2 h in DMEM low-glucose medium with 0.1% BSA. Cells were washed with Krebs Ringer phosphate Hepes (KRPH) buffer (140 mM NaCl, 5 mM KCl, 1.3 mM CaCl2, 1.2 mM KH2PO4, 2.5 mM MgSO4, 5 mM NaHCO3, 25 mM Hepes, and 0.1% BSA) and metabolically radiolabeled with 2 μCi/mL ^3^H-leucine (L-[3,4,5-^3^H(N)]-Leucine (Perkin-Elmer)) at 37°C for 15 mins. After incubation, cells were lysed in 0.5 N NaOH supplemented with 0.5% Triton X-100, and radioactive leucine uptake was measured using a Liquid Scintillation Counter and normalized to total protein content per well.

### Cell lines

Experiments were conducted in HEK 293T/17 cells (ATCC CRL-11268). This cell line was authenticated by ATCC based on its short tandem repeat profile and has recurrently tested negative for mycoplasma contamination.

### Plasmid transfection for biochemistry analysis

Transfection of GFP-tagged AnkB and mCherry-tagged Asct2 plasmids were conducted in HEK 293T/17 cells grown in 10 cm culture plates using the calcium phosphate transfection kit (Takara) and 5 µg of each plasmid DNA. Cell pellets were collected 48 hours after transfection for biochemistry analysis.

### Immunoblots

Protein homogenates from WAT, BAT, SKM and, liver or transfected cells were prepared in 1:7 (wt/vol) ratio of homogenization buffer (0.32M sucrose, 10mM HEPES, 1mM EDTA, 5mM N-ethylmeimide, 1mM sodium azide) supplemented with protease (#4693159001, Sigma) and phosphatase (#4906837001, Sigma) inhibitors cocktails. Total protein lysates were mixed at a 1:1 ratio with 5x PAGE buffer (5% SDS (wt/vol), 25% sucrose (wt/vol), 50mM Tris pH 8, 5mM EDTA, bromophenol blue) and heated for 15 min at 65°C. All lysates were resolved by SDS-PAGE on 3.5-17.5% acrylamide gradient gels in Fairbanks Running Buffer (40mM Tris pH 7.4, 20mM NaAc, 2mM EDTA, 0.2% SDS (wt/vol)). Proteins were transferred at 29V overnight onto 0.45 μm nitrocellulose membranes (#1620115, BioRad) at 4°C in methanol transfer buffer (25mM Tris, 1.92M Glycine, 20% methanol). Transfer efficiency was determined by Ponceau-S stain or Revert™ 520 or 700 Total Protein Stain (#926-10011, #926-11021, LI-COR). Membranes were blocked in TBS containing 5% non-fat milk for 1 hour at room temperature and incubated overnight with primary antibodies diluted in antibody buffer (TBS, 5% BSA, 0.1% Tween-20). After 3 washes in TBST (TBS, 0.1% Tween-20), membranes were incubated with secondary antibodies diluted in TBST containing 5% non-fat milk for two hours at room temperature. Membranes were washed 3x for 10 minutes with TBST and 2x for 5 minutes in TBS. Protein-antibody complexes were detected using the Odyssey® CLx or Odyssey® M imaging systems (LI-COR). Quantification of relative protein expression in western blot normalized to either an internal housekeeping protein control or total protein stain was conducted using the Empiria Studio Software (LI-COR).

### Immunoprecipitation

For immunoprecipitation experiments, total protein homogenates from transfected HEK 293T/17 cells were prepared in TBS containing 150 mM NaCl, 0.32 M sucrose, 2 mM EDTA, 1% Triton X-100, 0.5% NP-40, 0.1% SDS and compete protease inhibitor cocktail (Sigma). Cell lysates were incubated with rotation for 1 h at 4 °C and centrifuged at 100,000g for 30 min. Soluble fractions were collected and precleared by incubation with Protein A/G magnetic beads (#88802, Life Technologies) for 1 hour in the cold. Samples were subjected to immunoprecipitation in the presence of Protein G magnetic beads/antibody or Protein G magnetic beads/isotype control complexes overnight at 4 °C. Immunoprecipitation samples were resolved by SDS–PAGE and western blot and signal detected using the Odyssey® CLx or Odyssey® M imaging systems (LI-COR).

### Labeling and detection of biotinylated surface proteins

Cultured adipocytes were rinsed with ice-cold PBS containing 1mM MgCl2, 0.1mM CaCl2 (PBSCM) and incubated with 0.5 mg/ml Sulfo-NHS-SS-biotin (Biovision) for one 1 hour at 4°C. Reactive biotin was quenched by two consecutive 7-minute incubations with 20 mM glycine in PBSCM on ice. Cell lysates were prepared in RIPA buffer (50 mM Tris-HCl, 150 mM NaCl, 1.0% (v/v) NP-40, 0.5% (w/v) Sodium Deoxycholate, 1.0 mM EDTA, 0.1% (w/v) SDS and, 0.01% (w/v) sodium azide) supplemented with complete protease inhibitor cocktail (Sigma). Cell lysates were incubated with rotation for 1 hour at 4°C and centrifuged at 100,000 x g for 30 min. Soluble fractions were collected and incubated with high capacity NeutrAvidin™ agarose beads (#29200, Pierce) overnight at 4°C to capture biotinylated surface proteins. The beads were washed three times with TBST. Captured proteins were eluted in 5x-PAGE buffer and resolved by SDS-PAGE and western blot.

### Immunocytochemistry

Adipocytes grown in uncoated Ibidi dishes were washed with cold PBS, fixed with 4% PFA for 10 min, and permeabilized with 0.2% Triton-X100 in PBS for 10 min at room temperature. Cells were blocked with antibody buffer (4% BSA in PBS, 0.1% Triton X-100) for 30 min at room temperature and processed for staining as tissue sections. Secondary antibodies, diluted 1:300 in antibody buffer, were applied overnight at °4C. After three 5 min washes with PBS, adipocytes were incubated with BODIPY (1:1,000) and DAPI (1:1,000) diluted in PBS for one hour at room temperature. Cells were mounted for imaging with Prolong Gold Antifade Reagent (Life Technologies).

### Image acquisition and image analysis

Images of adipocytes stained were acquired using a Zeiss LSM780 confocal scope and 405-, 488-, 561- and 633-nm lasers using the Zeiss ZEN 2.3 SP1 FP1 (black) v.14.0.9.201 acquisition software. Single images and Z-stacks with optical sections of 1-μm intervals and tile scans were collected using the ×10 (0.4 numerical aperture (NA)) and ×40 oil (1.3 NA) objective lenses. Images were processed, and measurements taken and analyzed using NIH ImageJ software. Three-dimensional rendering of confocal Z-stacks was performed using Imaris (Bitplane).

### Statistical analysis

Sample size (n) was estimated using power analyses and expected effect sizes based on preliminary data in which we used similar methodologies, specimens, and reagents. We assumed a moderate effect size (f=0.25-0.4), an error probability of 0.05, and sufficient power (1-β=0.8). GraphPad Prism (GraphPad Software) was used for statistical analysis. Two groups of measurements were compared by unpaired Student’s t-test. Multiple groups were compared by one-way analysis of variance (ANOVA) followed by Tukey’s or Dunnett’s multiple comparisons test.

### Data and Resource Availability

The datasets generated during and/or analyzed during the current study are available from the corresponding author upon reasonable request. Gene expression profile data was deposited in GEO, access number GSE239665.

## RESULTS

### Elevated Levels of Circulating BCAA Precede the onset of Adiposity and Metabolic Dysregulation in Ankyrin-B Deficient Mice

We previously reported that whole body knock-in mice homozygous for the T2D-linked p.R1788W substitution within the unstructured C-terminal regulatory domain of the membrane scaffolding protein AnkB (*Ank2^R1788W/R1788W^*), herein referred as AnkB-RW mice, develop age-dependent obesity and glucose mishandling at 10-months of age (30). These metabolic deficits were also observed in young (3-month-old) AnkB-RW mice challenged with a high fat diet (30). As we previously showed, 3-month-old AnkB-RW mice have no significant differences in body weight (Fig 1A) or body composition (Fig 1B) when compared to littermate controls but exhibit increased insulin sensitivity (30). However, at this age AnkB-RW mice already display cell autonomous increases in white adipocyte size (30, 31), and in levels of fatty acid synthase (Fasn) and acyl-CoA synthetase (Acyl) in white adipose tissue (WAT) (Fig 1C, D). The enhanced fatty acid synthesis supports our previous findings showing that reduced AnkB levels in WAT leads to lipid accumulation in white adipocytes prior to the onset of obesity (30, 31).

**Figure 1.**
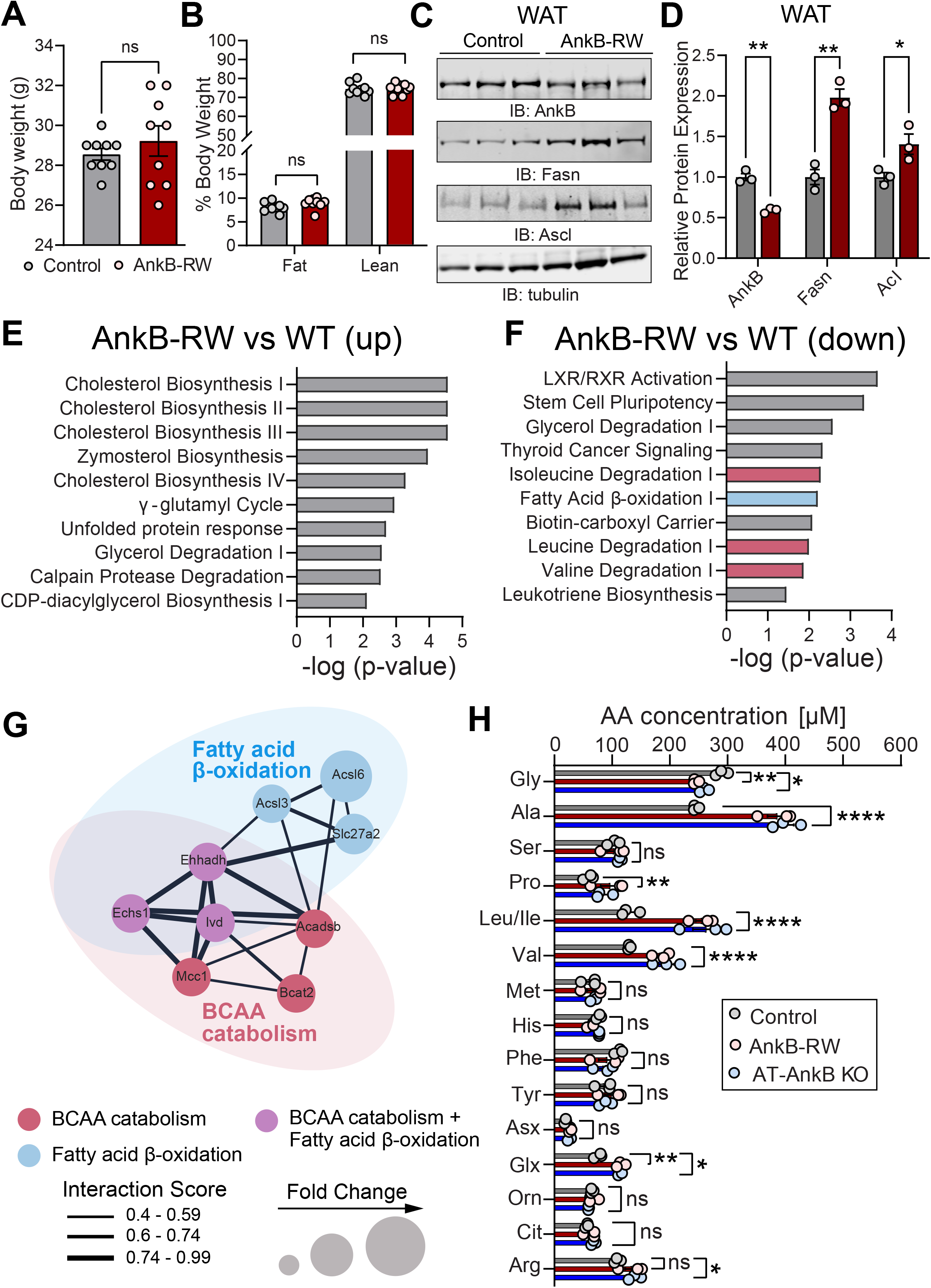
Reduced BCAA catabolism in WAT and elevated circulating BCAA precede the onset of obesity in ankyrin-B deficient mice. *A, B*: Body weight (*A*) and body composition (*B*) of 4-month-old control and AnkB-RW male mice. Data show mean ± SEM for n=9 animals per genotype. *C*: Representative immunoblots showing expression of AnkB, Fasn, and Ascl in WAT of lean 4-month-old control and AnkB-RW mice. *D:* quantification of protein levels in WAT normalized to tubulin expression. Data in *C* and *D* show mean ± SEM for three biological replicates per genotype representative of one of three independent experiments. *E, F*: Top functional pathways upregulated (*E*) and downregulated (*F*) in WAT from lean AnkB-RW compared to control mice. Data in *E* and *F* was derived from transcriptomic analysis from n=3 mice per genotype and analyzed using Ingenuity Pathway Analysis. *G*: Network analysis of interactions between genes involved in fatty acid oxidation (blue), BCAA catabolism (red), or both (purple) that are downregulated in WAT of AnkB-RW mice. Interaction scores were computed using STRING and visualized in Cytoscape. *H*: Levels of circulating amino acids in plasma of control, adipose tissue-specific AnkB knockout (AT-AnkB KO), and AnkB-RW mice. Data show mean ± SEM for n=3 animals per genotype. For all panels dots represent individual values. Data in A-C was analyzed by an unpaired t test. Data in H was analyzed by One-way ANOVA with Tukey’s post hoc analysis test for multiple comparisons. ****p <0.0001, **p <0.01, *p < 0.05, ^ns^p >0.05.

To further explore the mechanisms underlying the early remodeling of WAT in AnkB-RW mice, we conducted transcriptomic analysis from WAT of lean 3-month-old mice. Ingenuity Pathway Analysis identified an upregulation of cholesterol biosynthesis and cellular stress pathways in WAT of AnkB-RW mice (Fig 1E). Surprisingly, the analysis also revealed a significant downregulation of genes associated with the catabolism of the three BCAAs (Fig 1F). Gene interaction analysis further uncovered an overlapping network of downregulated genes involved in both fatty acid and BCAA oxidation in WAT of AnkB-RW mice (Fig 1G). Thus, coordinated impairments in lipid and BCAA utilization in WAT likely alter adipocyte energetics of AnkB-RW mice prior to the onset of obesity and systemic metabolic dysregulation, and further promote adipocyte hypertrophy.

Lower utilization of BCAA by peripheral tissues can result in their build up in the circulation (22). To assess whether the observed transcriptional changes in BCAA catabolic pathways in WAT alter systemic BCAA levels in AnkB-RW mice, we evaluated serum amino acids by targeted metabolomics. Levels of eight (Ser, Met, His, Phe, Tyr, Asx, Orn, Cit) out of the 16 amino acids measured were unchanged between AnkB-RW and control mice (Fig 1H). In contrast, there was a striking elevation of BCAAs, together with Ala, Pro, Glx, and Arg, and a reduction in Gly levels in serum of 3-month-old lean, non-IR, AnkB-RW mice (Fig 1H). The combined elevation of BCAA and reduction of Gly levels was also observed in serum from mice with conditional loss of AnkB in adipose tissue (*Ank2^f/f^;Adipoq* (AT-AnkB KO)) (Fig 1H). Interestingly, this serum amino acid signature has been recurrently detected in obese, IR humans compared to lean, insulin-sensitive individuals across many studies (3, 8, 22). Our findings that AnkB deficits lead to serum BCAA elevations in non-obese mice implicates AnkB in the regulation of BCAA flux and systemic BCAA levels. It also establishes AnkB-RW mice as a good model to investigate the systemic effects of early increases in circulating BCAAs in non-obese mice and its potential causal role in the pathogenesis of metabolic disorders.

### Impaired BCAA Catabolism in White Adipocytes in Ankyrin-B Deficient Mice Contribute to Elevated Levels of Circulating BCAA

Elevated levels of serum BCAA in obesity arise, in part, from their reduced oxidation due to coordinated transcriptional downregulation of BCAA catabolic enzymes (22). Prior studies suggest that reductions in BCAA catabolism in WAT directly contribute to the increases in plasma BCAA detected in obese and IR individuals, and in rodent models of these metabolic disease phenotypes (9, 28, 32). Therefore, we assessed transcriptional regulation of BCAA catabolism in different peripheral tissues of non-obese AnkB-RW mice to determine their potential contribution to the observed serum BCAA buildup. We confirmed that WAT from AnkB-RW mice expresses decreased levels of multiple BCAA oxidative enzymes, including Bcat2 and components of the mitochondrial branched-chain keto acid dehydrogenase (Bckdh) complex (Fig 2A, B). The Bckdh complex, which catalyzes the rate-limiting step of BCAA catabolism, is tightly regulated by inhibitory phosphorylation of the Bckdh e1 alpha subunit at serine 293 (pBckdh e1a^S293^) (33) (Fig 2A). We found that total protein levels of Bckdh e1a were slightly reduced in WAT of AnkB-RW mice concomitant with an increase in the relative expression of inactive pBckdh e1a^S293^(Fig 2C-D). In contrast, BCAA catabolism was not transcriptionally altered in liver, skeletal muscle, or brown adipose tissue (BAT) of AnkB-RW mice, except for decreased Bcat2 expression in BAT (Fig 3A-C), and no significant changes in protein expression or relative levels of inactive Bckdh e1a^S293^ were observed in these tissues (Fig. 3D-I). Collectively, these data suggest that increased plasma BCAA levels in ankyrin-B deficient mice are caused by impaired BCAA catabolism in WAT.

**Figure 2.**
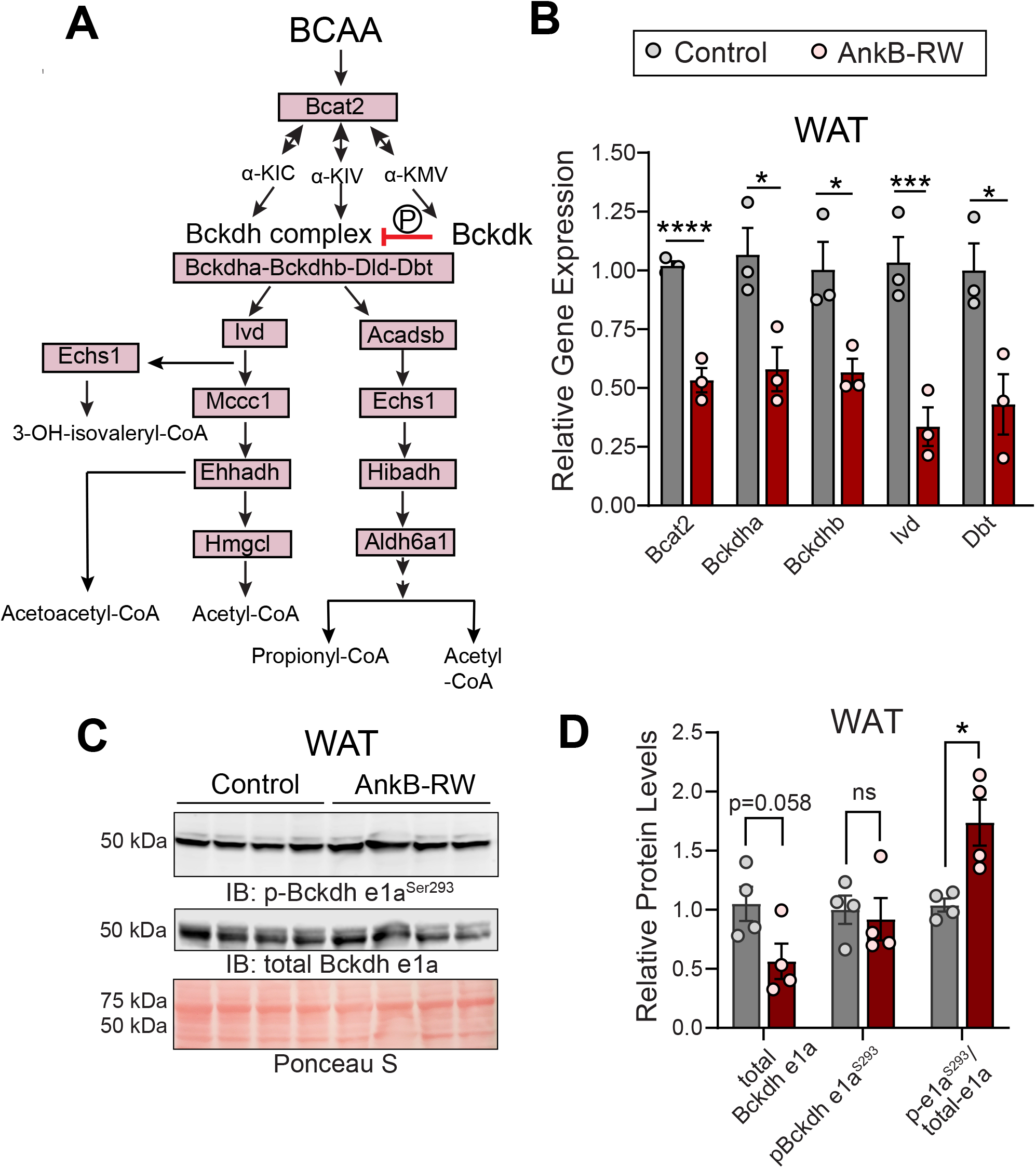
Elevated circulating BCAA levels in ankyrin-B deficient mice are associated with deficits in BCAA catabolism in WAT. *A*: Schematic of the major enzymes involved in BCAA catabolism. Upon cellular uptake, BCAAs are converted to branched chain α-ketoacids (BCKA) by mitochondrial Bcat2. BCKAs are further metabolized by the branched chain ketoacid dehydrogenase complex (Bckdh), which is deactivated by phosphorylation of the E1a subunit. BCAA catabolism contributes to the intracellular pool of acetoacetyl-Coa, acetyl-CoA, and propionyl-CoA. *B*: Relative gene expression of BCAA catabolic enzymes in WAT of lean 4-month-old control and AnkB-RW mice. *C*: Representative immunoblots showing expression of total Bckdhe1a and p-Bckdhe1a^S293^ in WAT of lean 4-month-old control and AnkB-RW mice. *D*: Relative expression of total Bckdhe1a and p-Bckdhe1a^S293^ levels normalized to total protein (Ponceau S), and of p-Bckdhe1a^S293^/Bckdhe1a ratio. For all panels dots represent individual values. Data was collected from n=3 mice per genotype and reported as mean ± SEM. Data was analyzed by an unpaired t test. ****p < 0.0001, ***p < 0.001, *p < 0.05.

**Figure 3.**
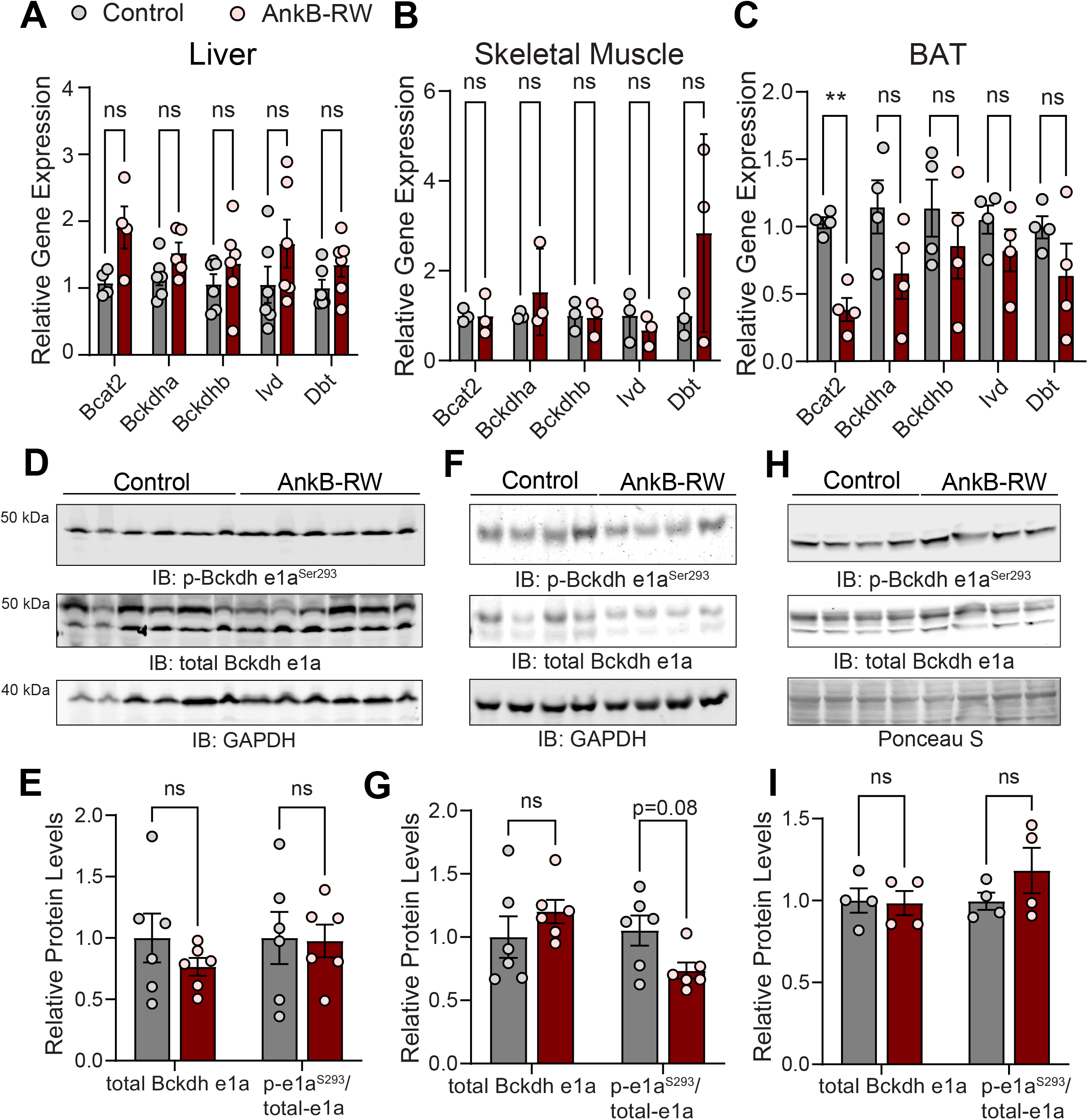
BCAA catabolism is not affected in liver, skeletal muscle, and brown adipose tissue of ankyrin-B deficient mice. *A-C*: Relative gene expression of BCAA catabolic enzymes in liver (*A*), skeletal muscle (*B*), and brown adipose tissue (BAT) (*C*) of lean 4-month-old control and AnkB-RW mice. *D-H*: Representative immunoblots showing expression of total Bckdhe1a and p-Bckdhe1a^S293^ in liver (*D*), skeletal muscle (*F*), and BAT (*H*) of lean 4-month-old control and AnkB-RW mice. *E-I*: Relative expression of total Bckdhe1a levels normalized to either GAPDH or total protein (Ponceau S), and of p-Bckdhe1a^S293^/Bckdhe1a ratio in liver (*E*), skeletal muscle (*G*), and BAT (*I*). For all panels dots represent individual values. Data was collected from n=4-6 mice per genotype and reported as mean ± SEM. Data was analyzed by an unpaired t test. ^ns^p >0.05.

### Ankyrin-B Deficiency Alters the Expression of BCAA Transporters in WAT

The amino acid transporter Lat1 (Slc7a5) forms a complex with Cd98 (Slc3a2) to facilitate BCAA uptake across the cell membrane against a glutamine gradient established by the Na^+^-dependent neutral amino acid exchanger Asct2 (Slc1a5) (34-36) (Fig 4A). To determine if the reduced BCAA catabolism observed in WAT of AnkB-RW mice is caused by changes in BCAA uptake, we evaluated the levels of BCAA transporters in WAT of these mice. We found that transcript and protein levels of the glutamine transporter Asct2 were decreased in WAT of AnkB-RW mice (Fig 4B-D), whereas CD98 and Lat1 expression were elevated (Fig 4B-C). In contrast, expression of Asct2, CD98, and Lat1 was unchanged in liver, skeletal muscle, and BAT of AnkB-RW mice (Supplementary Fig 1). Asct2 transcript and protein levels were also lower in WAT of 3-month-old AT-AnkB KO mice (Supplementary Fig 2). Remarkably, leucine uptake was reduced in primary white adipocytes from AnkB-RW mice despite the increases in CD98 and Lat1 (Fig 4E). Together, these observations suggest that reduction of Asct2 levels in WAT due to AnkB deficiency leads to impaired glutamine uptake and a shift in the chemical gradient that drives BCAA uptake by white adipocytes.

**Figure 4.**
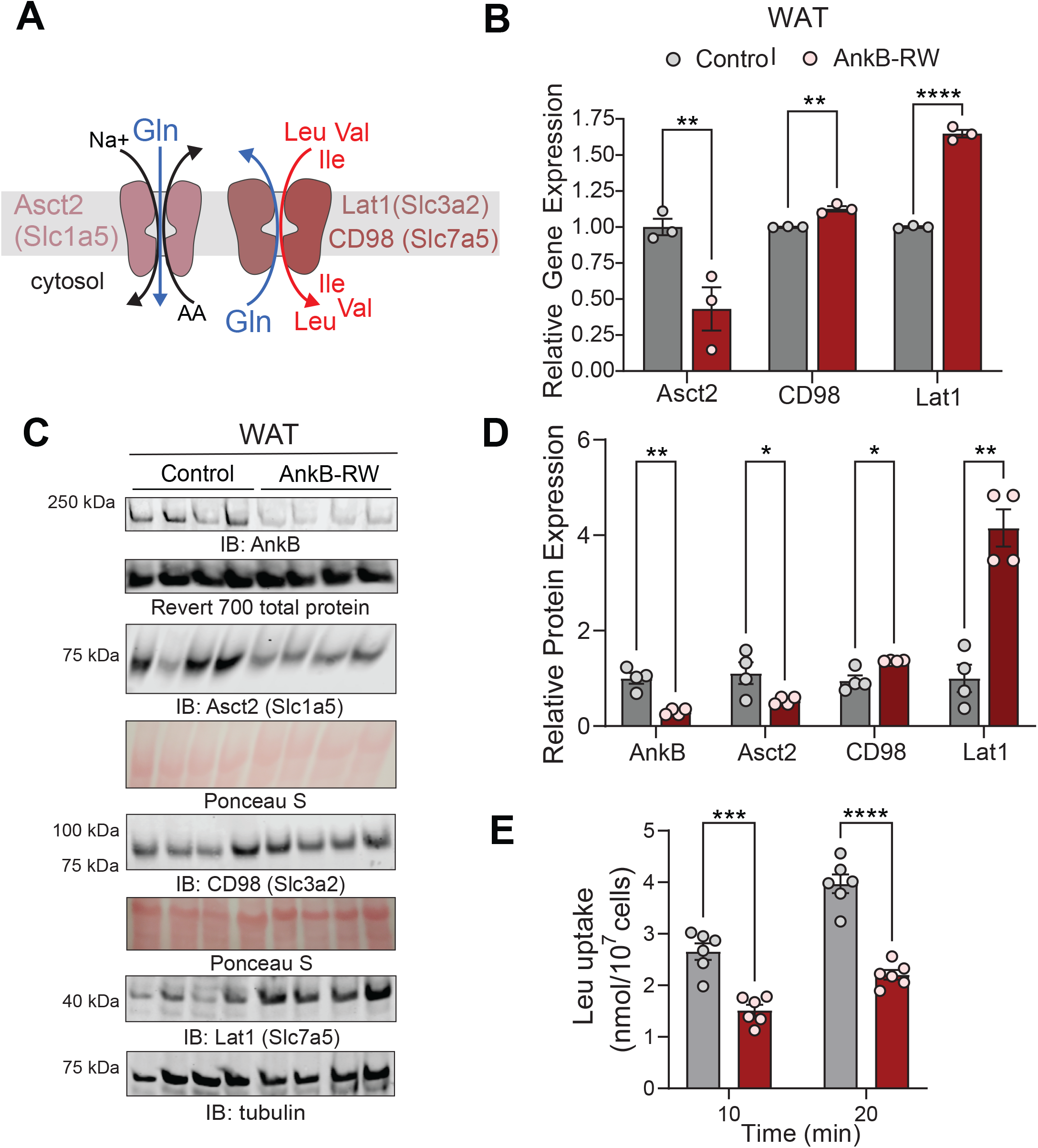
Ankyrin-B deficiency alters expression of BCAA transporters and BCAA uptake in WAT. *A*: Schematic of the major neutral amino acid transporters involved in cellular BCAA uptake. Asct2 (Slc1a5) transports glutamine into the cell establishing a chemical gradient that drives the uptake of BCAAs. In response to an increase in intracellular levels of glutamine, CD98 (Slc7a5), which exists as a heterodimer with Lat1 (Slc7a5), exports glutamine out of the cell while Lat1 (Slc7a5) simultaneously imports BCAAs. *B*: Relative gene expression of neutral amino acid transporters Asct2, CD98, and Lat1 in WAT of lean 4-month-old control and AnkB-RW mice. *C*: Representative immunoblots showing expression of AnkB and amino acid transporters in WAT of lean 4-month-old control mice. *D*: Relative expression of AnkB and amino acid transporters normalized to total protein (Revert 700, Ponceau S) or tubulin levels. *E*: Quantification of leucine uptake by differentiated primary white adipocytes. For all panels data show mean ± SEM with dots representing individual values. Data in *B* and *D* was collected from 3-4 mice per genotype. Data in *E* shows six biological replicates from two independent experiments. Data are reported as the mean ± SEM. Data was analyzed by an unpaired t test. ****p < 0.0001, ***p < 0.001, **p < 0.01, *p < 0.05.

### Ankyrin-B Interacts with and Modulates the Surface Levels of Asct2 in Adipocytes

AnkB promotes the localization and stability of diverse membrane proteins (37, 38). Upon AnkB deficits, its membrane-spanning partners are often mislocalized and dysregulated (30, 31, 39). The reduction in Asct2 levels detected in WAT and elevations of circulating glutamine (Fig 1H) in AnkB-RW mice led us to hypothesize that AnkB interacts with and promotes the surface localization of Asct2 in white adipocytes. To determine if AnkB binds Asct2, we co-expressed full-length (FL) GFP-tagged AnkB and mCherry-tagged Asct2 in HEK293 cells (Fig 5A). Evaluation of immunoprecipitation (IP) eluates confirmed that FL GFP-AnkB directly associates with mCherry-Asct2 (Fig 5B). We next sought to identify the sites of AnkB interaction with Asct2. Co-IP experiments revealed that Asct2 associates with the membrane binding domain of AnkB (MBD AnkB-GFP) containing the ankyrin (ANK) repeats, but not with its ZU5-Ct (ZU5-Ct AnkB-GFP) supermodule (Fig 5A, B).

**Figure 5.**
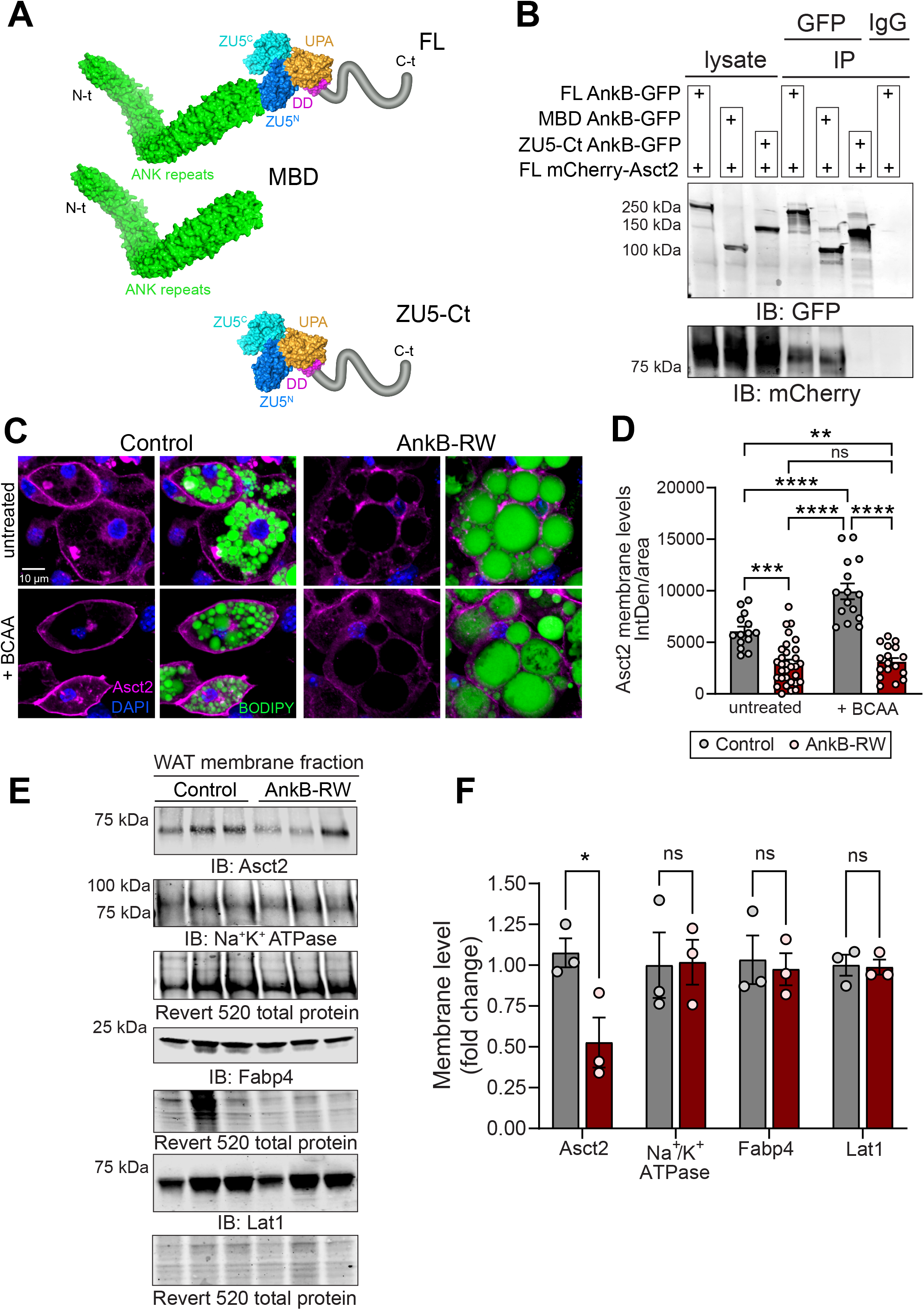
Ankyrin-B interacts with the BCAA transporter Asct2 and modulates its surface levels in adipocytes. *A*: Schematic depicting full-length (FL), membrane binding domain (MBD), and ZU5-C-terminal (ZU5-Ct) AnkB constructs used to evaluate AnkB interaction with Asct2. *B*: Representative immunoblots showing expression of GFP-AnkB proteins and mCherry-Asct2 in total lysates and immunoprecipitation (IP) eluates. *C*: Representative immunofluorescent images show Asct2 (magenta) distribution in differentiated white adipocytes at the basal state and upon BCAA treatment. BODIPY (green) labels lipid droplets. DAPI (bue) labels nuclei. *D*: Quantification of surface Asct2 in control (untreated n=14, BCAA-treated n=15) and AnkB-RW (untreated n=31, BCAA-treated n=17) white adipocytes. *E*: Representative immunoblots showing surface levels of indicated proteins in control and AnkB-RW cultured white adipocytes. *F*: Quantification of surface levels of Asct2, Na^+^/K^+^ ATPAse, Fabp4, and Lat1 normalized to total surface protein levels (Revert 520) collected from three independent experiments. For all panels data show mean ± SEM. Data in *D* was analyzed by One-way ANOVA with Tukey’s post hoc analysis test for multiple comparisons. Data in *F* was analyzed by an unpaired t test. ****p < 0.0001, ***p < 0.001, **p < 0.01, *p < 0.05, ^ns^p >0.05.

Finally, we asked whether AnkB deficiency perturbs Asct2 surface levels in white adipocytes. By evaluating Asct2 fluorescence signal, we found that AnkB-RW adipocytes had lower basal levels of membrane-associated Asct2 (Fig 5C, D). Remarkably, while treatment with BCAA increased surface localization of Asct2 by almost two-fold in control adipocytes, this effect was not observed in AnkB-RW cells (Fig 5C, D). In addition, using a biotinylation assay that captures surface proteins, we observed a marked reduction in surface levels of Asct2 in AnkB-RW white adipocytes relative to its membrane abundance in control adipocytes, while surface levels of other membrane proteins including Na^+^/K^+^ ATPase, Fabp4, and Lat1 remained unchanged (Fig 5E, F). Our findings identify AnkB as a specialized adaptor that is required for the proper localization of Asct2 at the cell surface of white adipocytes.

### Elevated Dietary BCAA Increases Fat Mass in Control Mice but does not Exacerbate Adiposity Phenotypes of Ankyrin-B Deficient Mice

Multiple studies suggest that elevated BCAA can fuel adipogenesis and promote adipose tissue expansion during obesity (40, 41) and contribute to IR (21, 22). AnkB-RW mice have elevated levels of plasma BCAA (Fig 1H) and reduced BCAA utilization by WAT (Fig 2B-D) prior to the onset of obesity (30). Thus, we reasoned that these mice would be more susceptible to metabolic dysfunction associated with exposure to BCAA-enriched diets. To test this prediction, we used a dietary approach to manipulate BCAA availability in which we randomly assigned control and AnkB-RW mice to one of four isocaloric diets with equal total protein content. Starting at weaning, mice were fed either a control diet (10% kcal fat) (CD), a CD supplemented with 1.5-fold BCAA (CD-BCAA), a high fat diet (HFD) (60% kcal fat), or a HFD supplemented with 1.5-fold BCAA (HFD-BCAA) for 12 weeks (Fig 6A). We confirmed that BCAA-enriched diets increased circulating BCAA levels in control mice, with the HFD-BCAA diet causing the most significant elevation (Supplementary Fig 3). HFD consumption also led to increases in serum BCAAs in control mice that were similar to the effects of the CD-BCAA formulation (Supplementary Fig 3). While CD-BCAA supplementation further increased serum levels of all BCAAs in AnkB-RW mice, HFD-BCAA feeding resulted in selective elevation of circulating valine (Supplementary Fig 3). Overall, BCAA-enriched diets did not alter physical activity, respiratory exchange ratio (RER) or energy expenditure (EE) of control or AnkB-RW mice (Supplementary Fig 4). In contrast, we found that BCAA supplementation, in both the context of CD or HFD, increased body weight and fat mass, and decreased lean mass in control mice (Fig 6B-D). Importantly, these changes were not due to increases in food consumption (Fig 6E). Control mice fed CD-BCAA or HFD-BCAA diets also exhibited hypertrophy of inguinal white adipocytes and increases in pro-inflammatory F4/80^+^ macrophages in WAT compared to mice of the same genotype respectively fed CD or HFD formulations (Fig 6F-H and Supplementary Fig 5). Consistent with our previous report (30), 4-month-old AnkB-RW mice were more susceptible to increases in body weight, fat mass, and adipocyte size associated with HFD intake (Fig 6B-H). Surprisingly, similar consumption of BCAA-enriched diets did not enhance adiposity of AnkB-RW mice (Fig 6B, C, E-G, Supplementary Fig 5). However, AnkB-RW mice fed HFD-BCAA showed the highest accumulation of F4/80^+^ macrophages in crown-like structures surrounding white adipocytes, suggesting increased inflammation (Fig 6H, Supplementary Fig 5).

**Figure 6.**
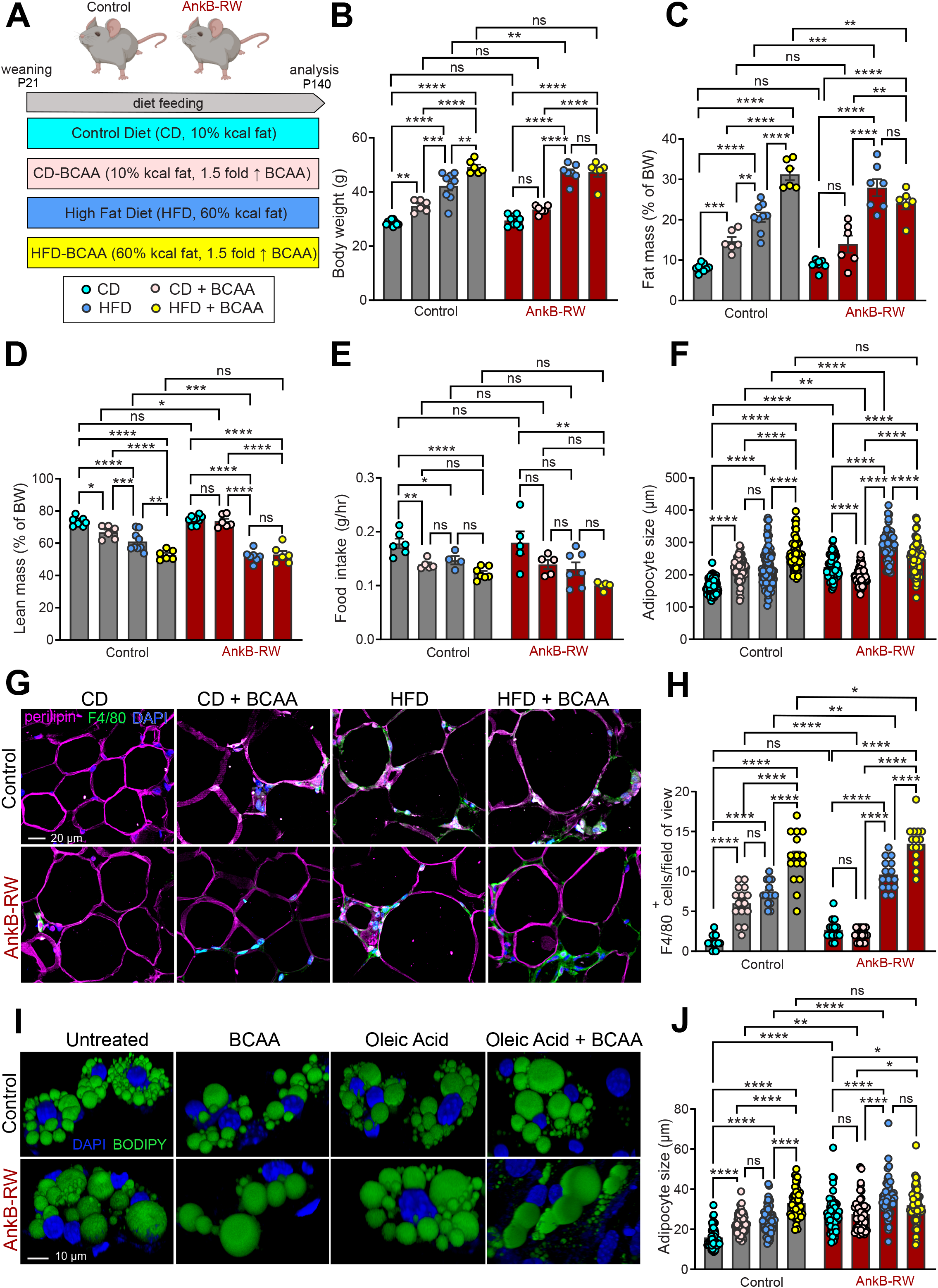
Exposure to elevated dietary BCAAs increases fat mass and adipocyte size in control mice but does not exacerbate adiposity phenotypes of ankyrin-B deficient mice. *A*: Schematic of experimental design for the diet feeding experiment. Control and AnkB-RW male mice (n= 7-10 mice per group) were fed either control diet (10% kcal fat) (CD), CD with 1.5-fold higher BCAA (CD-BCAA), high-fat diet (60% kcal fat) (HFD), or HFD with 1.5-fold higher BCAA (HFD-BCAA) content for 12 weeks. *B*: Body weight after 12 weeks of diet feeding. *C:* Fat mass as a percentage of body weight. *D*: Lean mass as a percentage of body weight. *E*: Food intake as grams/hour (g/hr). *F*: Adipocyte size quantified from WAT of 4-month-old control and AnkB-RW mice fed indicated diets for 12 weeks. *G*: Representative images of WAT sections from 4-month-old control and AnkB-RW mice fed indicated diets for 12 weeks. Sections were stained with perilipin to label adipocyte membranes, F4/80 to detect inflammatory macrophages. DAPI stains nuclei. Scale bar, 20 µm. *H*: Quantification of F4/80^+^ cells per field of view. *I*: Representative images of control and AnkB-RW adipocytes in culture media (untreated) or treated with either BCAA, oleic acid, and both oleic acid and BCAA. BODIPY labels lipid droplets. DAPI labels nuclei. Scale bar, 10 µm. *J*: Size of control and AnkB-RW differentiated white adipocytes. For all panels data show mean ± SEM. Data was analyzed by One-way ANOVA with Tukey’s post hoc analysis test for multiple comparisons. ****p < 0.0001, ***p < 0.001, **p < 0.01, *p < 0.05, ^ns^p >0.05.

BCAA catabolism has been proposed to fuel adipocyte differentiation and lipogenesis (40, 42). Given that AnkB-RW adipocytes exhibit deficient BCAA uptake and oxidation, we next compared their response to a BCAA-enriched differentiation cocktail. Using BODIPY-based quantification of lipids as a readout of adipogenesis, we found that a 1.5-fold increase in BCAA levels in the adipocyte differentiation media increased lipid droplet size in control adipocytes (Fig 6I, J). As expected, oleic acid treatment enhanced adipogenesis of control adipocytes relative to untreated cells, but its effects were similar to the BCAA treatment (Fig 6I, J). In contrast, control adipocytes exposed to a differentiation media supplemented with both BCAA and oleic acid exhibit the most prominent increases in lipid droplet size (Fig 6I, J), suggesting synergistic effects. We have previously shown that differentiated AnkB-RW adipocytes maintain elevated surface levels of the glucose transporter Glut4 due to deficits in AnkB-mediated coupling of Glut4 to the endocytic machinery, which results in increased glucose uptake and lipid accumulation (30, 31) (Fig 6I, J, untreated group). Lipid accumulation in AnkB-RW adipocytes was also enhanced by oleic acid (Fig 6I, J). However, BCAA supplementation, either alone or together with fatty acids, did not respectively increase lipid droplet size relative to untreated of oleic acid-treated AnkB-RW adipocytes (Fig 6I, J). Collectively, these findings indicate that AnkB-RW mice are protected from the obesogenic effects associated with increased BCAA utilization due to their impaired BCAA catabolism in WAT.

### Exposure to Elevated Dietary BCAA Worsens Glucose Regulation of Control and Ankyrin-B Deficient Mice

We next evaluated the effects of increased BCAA intake on systemic glucose regulation. Twelve weeks of BCAA-enriched diet feeding impaired glucose tolerance in control mice, demonstrated by elevations in fasting glucose levels and a delay in glucose clearance from the circulation following a glucose challenge (Fig 7A-C). As we reported, CD-fed AnkB-RW mice exhibit similar fasting serum glucose levels as control mice, but impaired glucose clearance during the glucose tolerance test (GTT) (Fig 7A-C) (30). In agreement with our prior findings, HFD-feeding most severely impaired fasting glycemia and glucose tolerance of AnkB-RW mice when compared to its effects on littermate controls (Fig 7A-C). Remarkably, consumption of BCAA-enriched diets exacerbated glucose handling deficits of AnkB-RW mice (Fig 7A), which contrasts with its negligible effects on adipogenesis (Fig. 6). HFD-BCAA feeding exerted the most deleterious impact on glucose tolerance across genotypes (Fig 7A-F), which supports the proposed negative effects of dual exposure to lipid and BCAA elevations (3, 50). While intake of BCAA-enriched diets or HFD promoted glucose-stimulated insulin secretion in 4-month-old control mice (Fig 7D), these stimulatory effects were blunted in AnkB-RW mice, especially those exposed to HFD-BCAA feeding, which exhibited further reductions in insulin secretion (Fig 7D). Lastly, we found that dietary BCAA supplementation worsened insulin responses in control mice during an insulin tolerance test (ITT), whereas these deleterious effects were only triggered in AnkB-RW mice exposed to dual BCAA and fat-rich diets (Fig 7E, F). Combined, these results suggest that synergistic exposure to chronic elevation in circulating lipids, BCAAs and other amino acids, potentiate insulin secretion deficits of AnkB-RW mice (30). These findings also raise the possibility that the obesogenic effects we found to be associated with increased BCAA consumption in control mice significantly contribute to systemic glucose dysregulation.

**Figure 7.**
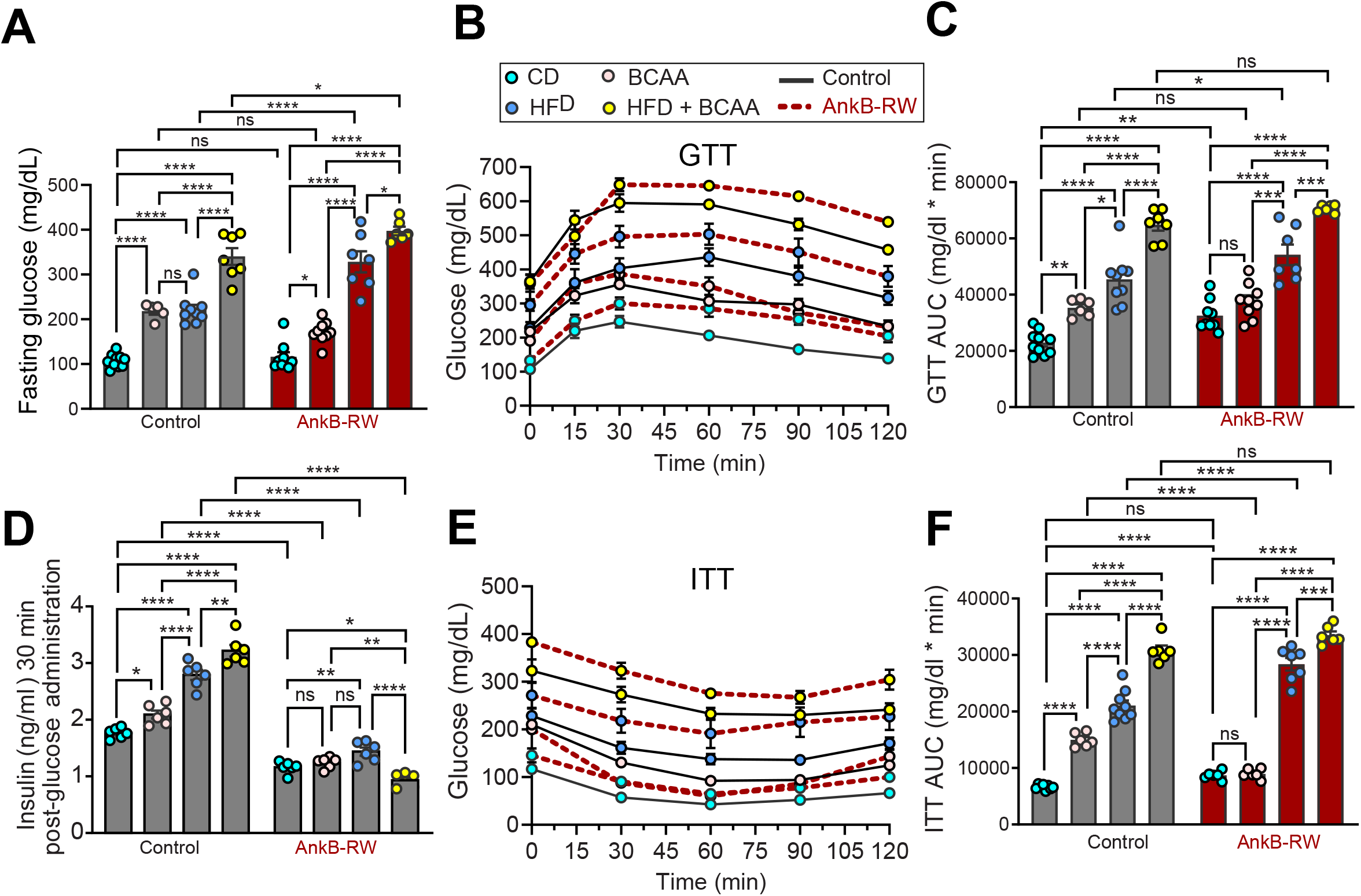
Exposure to elevated dietary BCAA worsens glucose regulation of control and ankyrin-B deficient mice. *A*: Fasting glucose levels of control and AnkB-RW mice fed indicated diets for 12 weeks. *B*: Glucose tolerance test (GTT) performed after 12 weeks of diet treatment. *C*: Area under the curve (AUC) showing glucose level changes during the GTT. *D*: Insulin levels 30 minutes post glucose administration. *E*: Insulin tolerance test (ITT) performed after 12 weeks of diet treatment. *F*: AUC showing glucose level changes during the ITT. Data was acquired from n= 4-10 mice per group. For all panels data show mean ± SEM. Data was analyzed by One-way ANOVA with Tukey’s post hoc analysis test for multiple comparisons. ****p < 0.0001, ***p < 0.001, **p < 0.01, *p < 0.05, ^ns^p >0.05.

## DISCUSSION

In this study, we identify a cell-autonomous mechanism that causes impairment of BCAA utilization in mice with targeted knockout of ankyrin-B in adipose tissue (AT-AnkB KO) or bearing the p.R1788W ankyrin-B (AnkB-RW) variant associated with increased risk for T2D and cardio-metabolic syndrome (30, 31, 43). We found that ankyrin-B mutant mice exhibited significant elevations in serum BCAAs prior to the onset of obesity and systemic lipotoxicity. Our results reveal that BCAA increases in the circulation of mice with ankyrin-B deficiency are due to selective impairment of BCAA catabolism in WAT. We demonstrated that WAT of non-obese, non-IR AnkB-RW mice exhibit reduced transcriptional and protein expression of BCAA catabolic enzymes concomitant with increased inactivation of the Bckdh complex, the rate limiting enzyme of BCAA catabolism. In contrast, BCAA catabolism did not change in liver, skeletal muscle, and BAT. We found that white adipocytes of AnkB-RW mice also have changes in expression of the neutral amino acid transporters Asct2, Lat1, and CD98 and reduced BCAA uptake. Mechanistically, we uncovered that ankyrin-B binds Asct2 and regulates its surface abundance in WAT. Therefore, we propose that ankyrin-B also modulates BCAA utilization by directly promoting BCAA uptake into white adipocytes. Finally, we show that consumption of BCAA-enriched diets increased body weight, fat mass and adipocyte size in control mice, but not in AnkB-RW mice, providing evidence that BCAA utilization by WAT drives adipogenesis and WAT expansion during obesity.

We found reduced BCAA catabolism in WAT of two non-obese mouse models of ankyrin-B deficiency (AT-AnkB KO and AnkB-RW mice), which exhibit increased glucose uptake by white adipocytes (30, 31). These findings are in line with a report of similar effects in mice overexpressing Glut4 in WAT (9). Thus, it is possible that increases in glucose disposal in WAT diminish the need to use BCAAs as substrates for fatty acid synthesis and contribute to systemic BCAA elevations. Our results showing that ankyrin-B binds Asct2 and controls its surface abundance in white adipocytes suggest it dually regulates BCAA utilization in WAT by modulating its rate of catabolism, secondary to cellular glucose availability, and by directly promoting BCAA uptake.

Utilization of BCAAs by WAT has been proposed to fuel *de novo* fatty acid synthesis and promote the initial stages of adipocyte differentiation (40, 42). Interestingly, blocking Asct2 reduces adipogenesis in 3T3-L1 adipocytes (44). Recent studies also suggest that enhancing BCAA catabolism in WAT of Bckdk knockout mice cell-autonomously promotes obesogenic expansion of subcutaneous fat (sWAT) (41). Conversely, blocking adipose BCAA catabolism, by ablation of either Bcat2 or Bckdha expression, attenuates diet-induced obesity (41). These beneficial effects are attributed, in part, to increases in sWAT browning and thermogenesis (45). Furthermore, lean mice lacking the mitochondrial phosphatase 2C (PP2Cm), which modestly impairs BCAA catabolism in different tissues simultaneously, showed lower body weight (46). We also found that BCAA supplementation to control mice, alone or together with fatty acids, increases lipid accumulation and fat mass *in vivo* and in cultured adipocytes. In contrast, AnkB-RW mice, which exhibit reduced BCAA uptake and catabolism in WAT, were protected from the adipogenic effects associated with increased BCAA availability. Our studies do not exclude the possibility that impaired BCAA catabolism in WAT of ankyrin-B deficient mice may in parallel exert beneficial effects on systemic energy balance by promoting BAT function. However, we found that AnkB-RW mice are susceptible to metabolic deficits associated with consumption of fat and BCAA-enriched diets likely through synergistic, systemic deleterious effects of elevated serum BCAAs and targeted-effects of ankyrin-B deficiency in peripheral tissues.

Multiple studies strongly support a causative role of elevations in circulating BCAAs, secondary to their reduced catabolism, in the development of IR. Promoting systemic BCAA oxidation through administration of the BCKD kinase inhibitor 3,6-dichlorobenzo(b)thiophene-2-carboxylic acid (BT2) lowers plasma BCAAs, improves glucose metabolism, and lessens IR in rodent models of obesity (25, 47). Likewise, boosting BCAA catabolism by treatment with sodium phenylbutyrate (NaPB), an analogue of BT2 that inhibits BCKD kinase at the same allosteric site, reduces both plasma BCAA and glucose levels, and improves peripheral insulin sensitivity in patients with T2D (48). However, these studies did not assess whether pharmacological treatment with BT2 or NaPB boost BCAA oxidation in WAT, and the direct contribution of elevated BCAA utilization by WAT to systemic improvement in plasma BCAA levels and glucose homeostasis.

Dietary BCAA restriction improves glucose tolerance and slows fat mass gain of healthy mice (12) and promotes fat mass loss in obese mice (49). Reduced BCAA intake also increases insulin sensitivity of Zucker-fatty rats (11). Conversely, the effects of BCAA-enriched diets on body weight, insulin sensitivity, and systemic metabolic health differ depending on the diet formulation, experimental design, and the rodent model. For instance, Wistar rats fed HFD-BCAA exhibit worse insulin sensitivity, despite lower body weight, than HFD-fed rats, whereas consumption of a BCAA-enriched chow diet has no effects on body weight or insulin sensitivity (3). We found that feeding C57BL6/J mice a low-fat diet supplemented with BCAAs (CD-BCAA) for 12 weeks increases fat mass and adipocyte size. Similar increases in body weight and adiposity were caused by long-term feeding of a low-fat diet supplemented with BCAAs to C57BL6/J mice, although, unlike in our study, these effects were accompanied by hyperphagia (13). We also show that feeding lean C57BL6/J mice diet pellets enriched in both BCAA and fats (HFD-BCAA), increases plasma BCAAs, and worsens obesity and glucose homeostasis. In contrast, a study in which C57BL6/J mice received BCAA supplementation in water simultaneously with HFD feeding did not observe additive deleterious effects on body weight or glucose regulation compared to the HFD-fed group (14). Differences in the route used for BCAA supplementation might partially account for the discrepancies in the systemics effects. Noteworthy, in that study BCAA supplementation to HFD-fed mice did not alter circulating levels of leucine and isoleucine but slightly increased plasma valine (14). Interestingly, feeding mice a valine enriched HFD drives adiposity and impairs glucose tolerance and insulin sensitivity (50). These negative effects have been attributed to an accumulation of the valine-derived metabolite 3-hydroxyisobutyrate (3-HIB) in plasma, and the concomitant glucotoxicity in skeletal muscle, which compromises insulin signaling (50). Thus, it would be important to define a threshold at which increases in BCAA, and specifically valine in the circulation promotes toxic 3-HIB accumulation and its deleterious effects.

Lastly, modulation of brain insulin via selective induction of hypothalamic insulin signaling in rats or genetic modulation of brain insulin receptors in mice indicate that the brain regulates BCAA metabolism by inducing hepatic BCKDH (51, 52). Additionally, impaired neutral amino acid transport in the brain caused by deficits in Lat1, which has been linked to autism, in either the blood brain barrier (53) or neurons (54), shifts BCAA levels and leads to neurological deficits. Thus, given the critical roles that ankyrin-B plays in the central nervous system and that its deficiency in brain is associated with neurodevelopmental disorders, including autism (37, 39, 55, 56), it would be informative to investigate whether AnkB deficits in the brain also contribute to the modulation of systemic or local BCAA levels.

## Acknowledgements

The authors would like to thank Dr. Olga R. Ilkayeva (Metabolomics Core Laboratory at Duke Molecular Physiology Institute) and Dr. Holly K. Dressman (Duke Sequencing and Genomic Technologies Shared Resource), both at Duke University School of Medicine, Durham, USA for their respective technical assistance with the metabolomic and gene expression analysis experiments. The authors would also like to thank Saame Shaikh and Traci Davis (Genomics and Energy Metabolism Core, University of North Carolina Nutrition and Obesity Research Center, Chapel Hill, USA) for their assistance with body composition and indirect calorimetry measurements.

## Authors Contributions

A.M.A., J.T., and D.N.L conceptualized and designed the study. A.M.A., J.T., I.F., B.T.T. and D.N.L conducted the experiments, data analyses and interpretation. A.M.A., J.T., I.F., and D.N.L wrote the manuscript. D.N.L. supervised the project and secured funding. D.N.L. is the guarantor of this work and, as such, had full access to all the data in the study and takes responsibility for the integrity of the data and the accuracy of the data analysis.

## Figure legends

**Figure S1:**
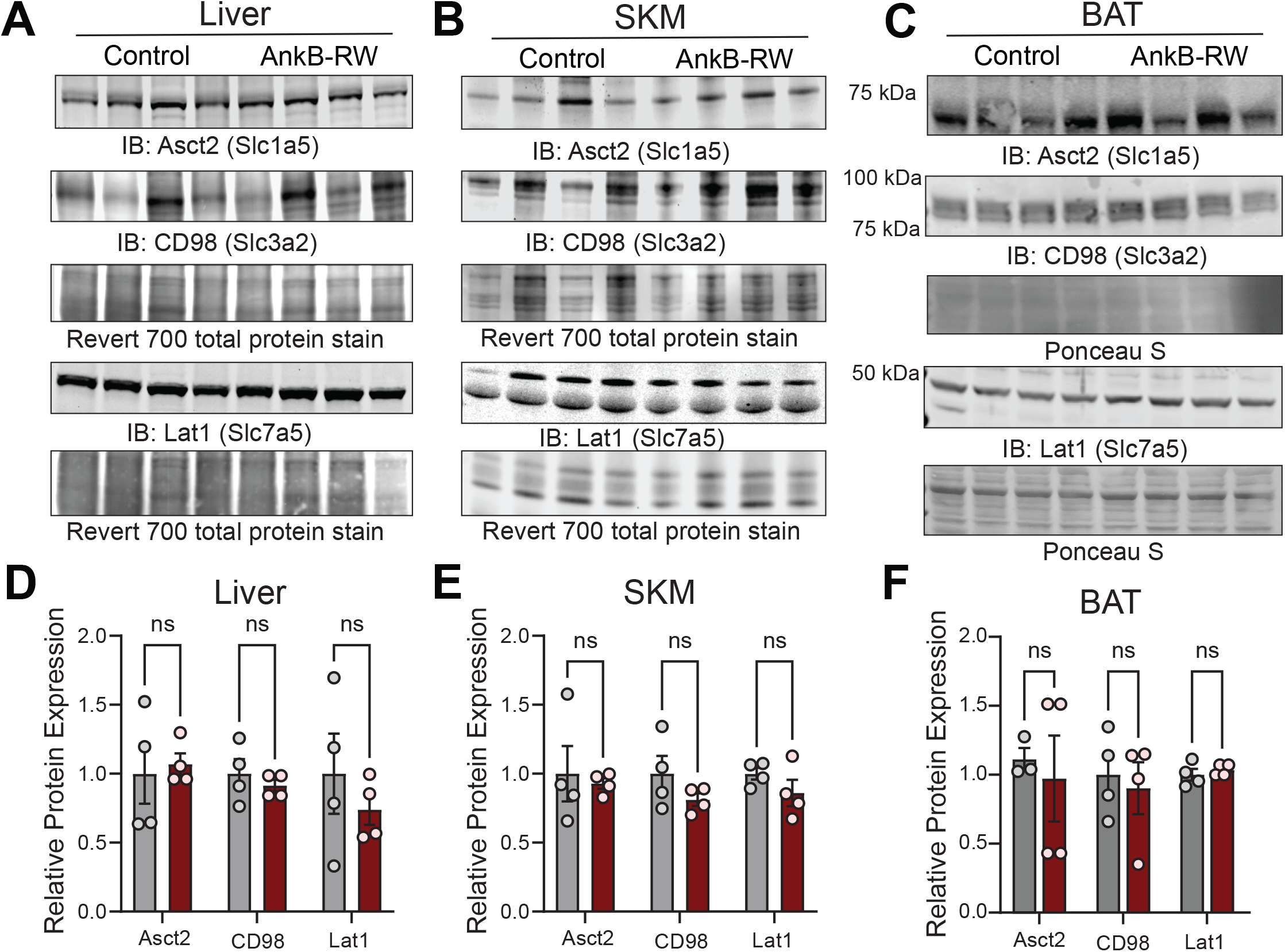
Ankyrin-B deficiency does not alter levels of BCAA transporters in liver, skeletal muscle, and BAT. *A-C*: Representative immunoblots showing expression of neutral amino acid transporters Asct2, CD98 and Lat1 in liver *(A)*, skeletal muscle *(B)*, and BAT *(C)* of lean 4-month-old control and AnkB-RW mice. *D-F*: Relative expression of indicated amino acid transporters normalized to total protein (Revert 700, Ponceau S). Data was collected from 4 mice per genotype. For all panels data show mean ± SEM with dots representing individual values. Data was analyzed by an unpaired t test. ^ns^p >0.05.

**Figure S2:**
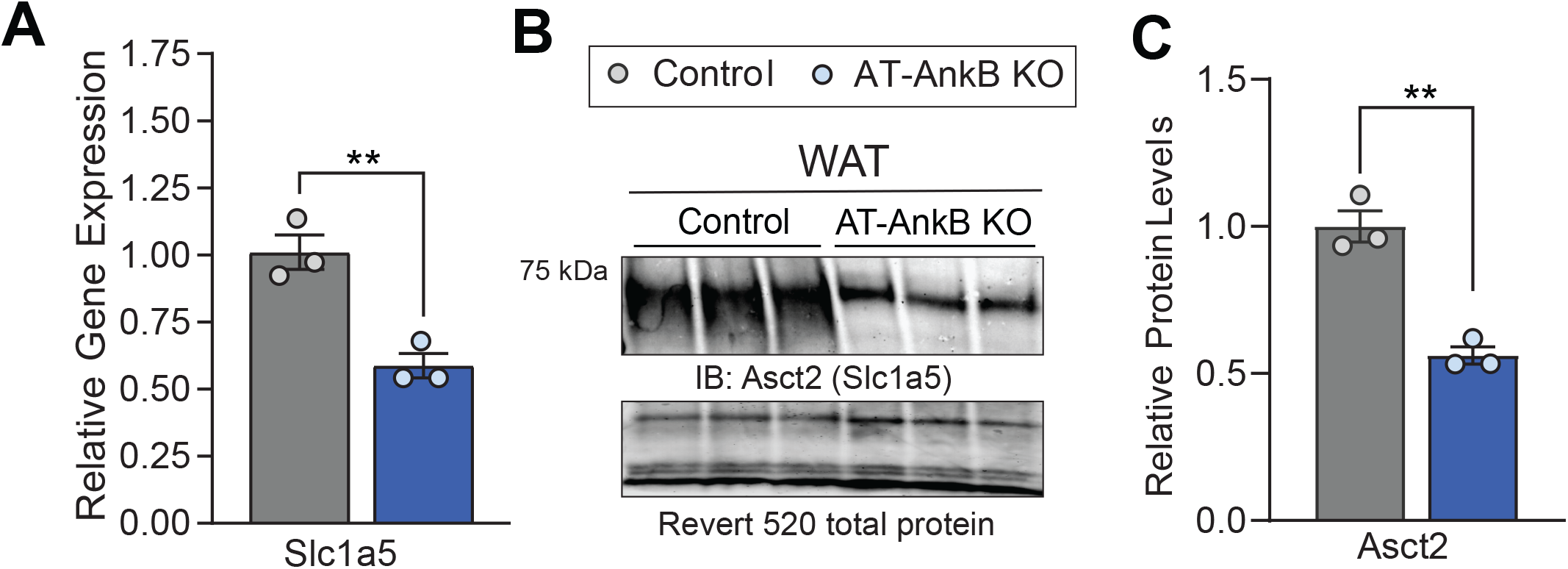
Lean AT-AnkB KO mice exhibit reduced Asct2 expression in WAT. *A*: Relative gene expression of Asct2 in WAT of lean 3-month-old control and AT-AnkB KO mice. *B*: Representative immunoblots showing expression of Asct2 in WAT of lean 3-month-old mice. *C*: Relative Asct2 expression normalized to total protein (Revert 520) levels. For all panels data show mean ± SEM with dots representing individual values. Data was collected from 3 mice per genotype. Data was analyzed by an unpaired t test. **p < 0.01.

**Figure S3:**
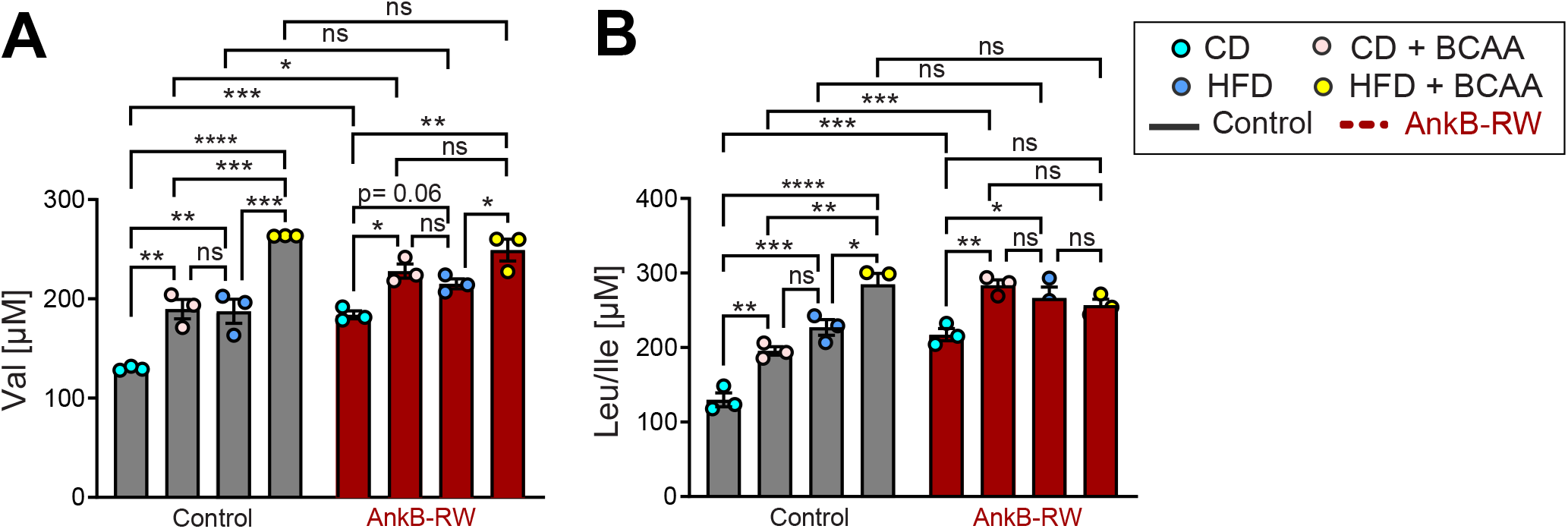
Plasma BCAA levels of control and AnkB-RW mice exposed to BCAA-enriched diets. *A, B*: Valine (Val) (*A*) and leucine/isoleucine (Leu/Ile) (*B*) levels in plasma of control and AnkB-RW mice (n=3 per group) fed indicated diets for 12 weeks. For all panels data show mean ± SEM. Data was analyzed by One-way ANOVA with Tukey’s post hoc analysis test for multiple comparisons. ****p < 0.0001, ***p < 0.001, **p < 0.01, *p < 0.05, ^ns^p >0.05.

**Figure S4:**
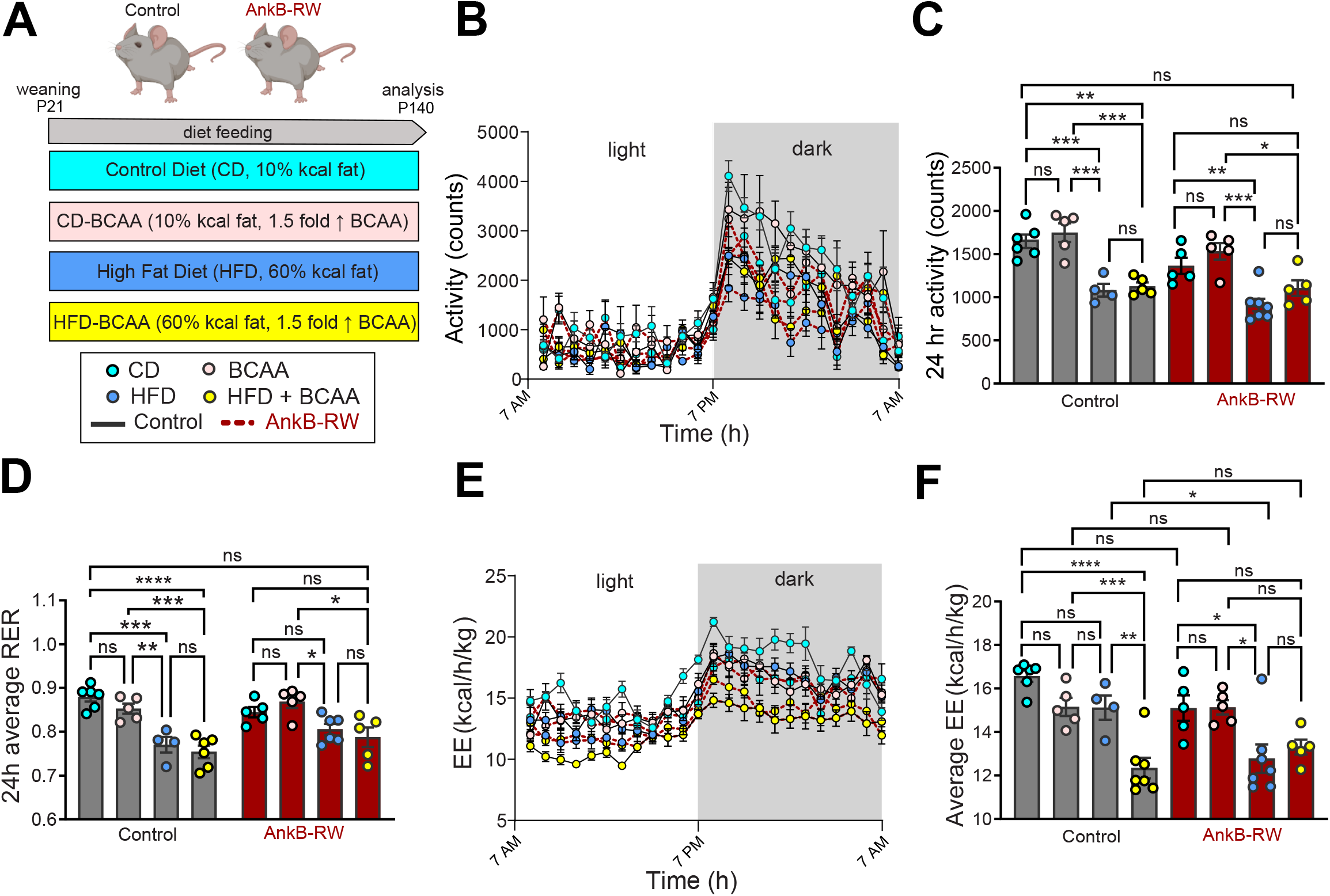
Indirect calorimetry analysis of control and AnkB-RW mice exposed to BCAA-enriched diets. *A*: Schematic of experimental design for the diet feeding experiment. Control and AnkB-RW male mice (n= 7-10 mice per group) were fed either control diet (10% kcal fat) (CD), CD with 1.5-fold higher BCAA (CD-BCAA), high-fat diet (60% kcal fat) (HFD), or HFD with 1.5-fold higher BCAA (HFD-BCAA) content for 12 weeks. *B*: Activity during light and dark cycles after 12 weeks of diet feeding. *C:* Average activity during a 24-hour period. *D*: Average RER during a 24-hour period. *E*: energy expenditure (EE) during light and dark cycles after 12 weeks of diet feeding. *F*: Average EE during a 24-hour period. For all panels data show mean ± SEM. Data was analyzed by One-way ANOVA with Tukey’s post hoc analysis test for multiple comparisons. ****p < 0.0001, ***p < 0.001, **p < 0.01, *p < 0.05, ^ns^p >0.05.

**Figure S5:**
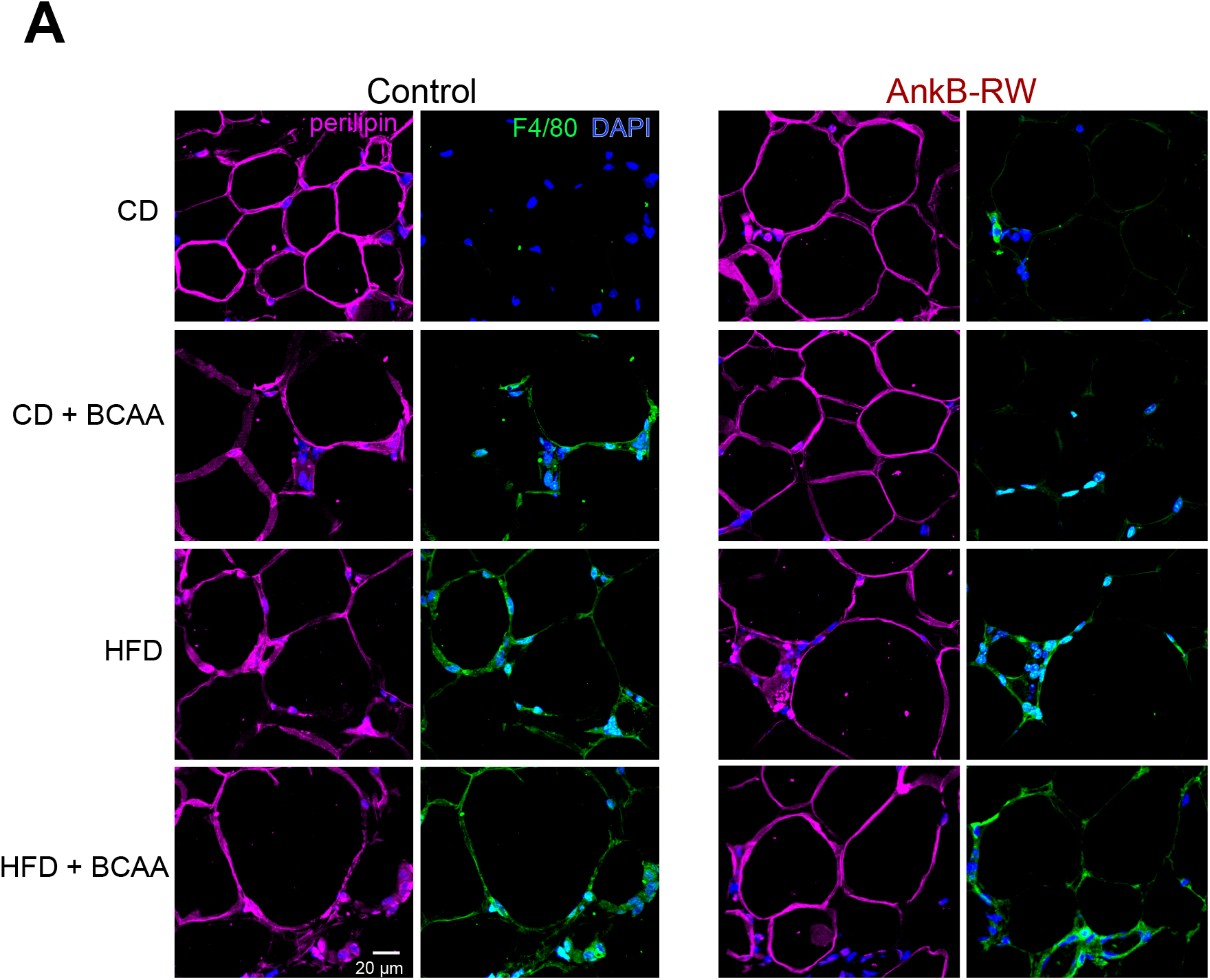
Effect of different diets on white adipocyte size and inflammation. *A*: Representative images of WAT sections from 4-month-old control and AnkB-RW mice fed indicated diets for 12 weeks. Sections were stained with perilipin (magenta) to label adipocyte membranes, F4/80 (green) to detect inflammatory macrophages. DAPI (blue) stains nuclei. Scale bar, 20 µm.

**Supplementary Table 1:** Caloric and nutritional composition of diets used in the study. All diet formulations were provided by Research Diet.

| Diet Component | HFD<br>D12492 |  | HFD-BCAA<br>D17050303 |  | CD<br>D12450K |  | CD-BCAA<br>D17011602 |  |
| --- | --- | --- | --- | --- | --- | --- | --- | --- |
| % | gm | kcal | gm | kcal | gm | kcal | gm | kcal |
| Protein | 26.2 | 20.0 | 29.6 | 23.0 | 19.2 | 20.0 | 22.0 | 23.0 |
| Carbohydrate | 26.3 | 20.1 | 25.0 | 19.4 | 67.3 | 70.0 | 64.7 | 67.4 |
| Fat | 34.9 | 59.9 | 33.1 | 57.7 | 4.3 | 10.0 | 4.1 | 9.6 |
| Total |  | 100 |  | 100 |  | 100 |  | 100 |
| kcal/gm | 5.24 |  | 5.16 |  | 3.85 |  | 3.84 |  |
| **Ingredient** | **gm** | **kcal** | **gm** | **kcal** | **gm** | **kcal** | **gm** | **kcal** |
| Casein, 80 Mesh | 200 | 800 | 100 | 400 | 200 | 800 | 100 | 400 |
| L-Cystine | 3 | 12 | 3.35 | 13 | 3 | 12 | 3.35 | 13 |
| **Isoleucine** | 0 | 0 | 16.8 | 67 | 0 | 0 | 16.8 | 67 |
| **Leucine** | 0 | 0 | 30.96 | 124 | 0 | 0 | 30.96 | 124 |
| Lysine | 0 | 0 | 6.5 | 26 | 0 | 0 | 6.5 | 26 |
| Methionine | 0 | 0 | 2.3 | 9 | 0 | 0 | 2.3 | 9 |
| Phenylalanine | 0 | 0 | 4.4 | 18 | 0 | 0 | 4.4 | 18 |
| Threonine | 0 | 0 | 3.35 | 13 | 0 | 0 | 3.35 | 13 |
| Tryptophan | 0 | 0 | 1.05 | 4 | 0 | 0 | 1.05 | 4 |
| **Valine** | 0 | 0 | 20 | 80 | 0 | 0 | 20 | 80 |
| Histidine | 0 | 0 | 2.3 | 9 | 0 | 0 | 2.3 | 9 |
| Alanine | 0 | 0 | 2.3 | 9 | 0 | 0 | 2.3 | 9 |
| Arginine | 0 | 0 | 3.15 | 13 | 0 | 0 | 3.15 | 13 |
| Aspartic Acid | 0 | 0 | 5.7 | 23 | 0 | 0 | 5.7 | 23 |
| Glutamic Acid | 0 | 0 | 18.4 | 74 | 0 | 0 | 18.4 | 74 |
| Glycine | 0 | 0 | 1.6 | 6 | 0 | 0 | 1.6 | 6 |
| Proline | 0 | 0 | 10.3 | 41 | 0 | 0 | 10.3 | 41 |
| Serine | 0 | 0 | 4.75 | 19 | 0 | 0 | 4.75 | 19 |
| Tyrosine | 0 | 0 | 4.65 | 19 | 0 | 0 | 4.65 | 19 |
| Corn Starch | 0 | 0 | 0 | 0 | 550 | 2200 | 550 | 2200 |
| Maltodextrin 10 | 125 | 500 | 125 | 500 | 150 | 600 | 150 | 600 |
| Sucrose | 68.8 | 275 | 68.8 | 275 | 0 | 0 | 0 | 0 |
| Cellulose, BW200 | 50 | 0 | 50 | 0 | 50 | 0 | 50 | 0 |
| Soybean Oil | 25 | 225 | 25 | 225 | 25 | 225 | 25 | 225 |
| Lard | 245 | 2205 | 245 | 2205 | 20 | 180 | 20 | 180 |
| Mineral Mix S10026 | 10 | 0 | 10 | 0 | 10 | 0 | 10 | 0 |
| DiCalcium Phosphate | 13 | 0 | 13 | 0 | 13 | 0 | 13 | 0 |
| Calcium Carbonate | 5.5 | 0 | 5.5 | 0 | 5.5 | 0 | 5.5 | 0 |
| Potassium Citrate, 1 H<sub>2</sub>O | 16.5 | 0 | 16.5 | 0 | 16.5 | 0 | 16.5 | 0 |
| Sodium BiCarbonate | 0 | 0 | 3.5 | 0 | 0 | 0 | 3.5 | 0 |
| Vitamin Mix V10001 | 10 | 40 | 10 | 40 | 10 | 40 | 10 | 40 |
| Choline Bitartrate | 2 | 0 | 2 | 0 | 2 | 0 | 2 | 0 |
| FD&C Yellow Dye #5 | 0 | 0 | 0.05 | 0 | 0 | 0 | 0.025 | 0 |
| FD&C Red Dye #40 | 0 | 0 | 0 | 0 | 0.025 | 0 | 0.025 | 0 |
| FD&C Blue Dye #1 | 0.05 | 0 | 0 | 0 | 0.025 | 0 | 0 | 0 |
| **Total** | **773.85** | **4057** | **816.21** | **4213** | **1055.05** | **4057** | **1097.41** | **4212** |

